# Sustained exposure to trypsin causes cells to transition into a state of reversible stemness that is amenable to transdifferentiation

**DOI:** 10.1101/679928

**Authors:** Maryada Sharma, Rajendra Kumar, Swati Sharma, Beena Thomas, Gargi Kapatia, Gurpreet Singh, Amanjeet Bal, Jagat Ram, Manoj Bhasin, Purnananda Guptasarma, Manni Luthra-Guptasarma

**Author notes:** Address correspondence to Dr. Maryada Sharma OR Prof. Manni Luthra-Guptasarma. Current affiliation: Department of Otolaryngology and Head and Neck Surgery, Postgraduate Institute of Medical Education and Research (PGIMER), Sector-12, Chandigarh 160012, India (Maryada Sharma).

## Abstract

During cell culture, trypsin, a serine protease, is applied to cells for 5-10 minutes to separate them from each other and from the underlying substratum so that they can be transferred to a different vessel, for re-plating, after growth medium containing 10 % serum has been added to the cells, in a well-known technique known as ‘passaging’. The serum in the growth medium contains alpha-1 antitrypsin, which is a potent inhibitor of trypsin, elastase and other serine proteases. Although what is used is bovine serum in which levels of proteins could be different from levels seen in humans, normal human serum contains A1AT (> 1 mg/ml; > ∼18 µmol/L) as well as trypsin itself (< 460 ng/ml, or ∼0.02 µmol/L), with the former in a ∼900-fold molar excess over the latter. Thus, it may be assumed there is also enough A1AT in the bovine serum added during passaging, to neutralize the trypsin (∼100 μM) present in the small volume of trypsin-EDTA solution used to separate cells. What are the consequences of not adding serum, when growth medium is added, or of maintaining cells for a few tens of hours in the presence of trypsin, in a serum-free growth medium? What does such sustained exposure to trypsin during cell culture do to cells? More generally, what are the responses of cells within an organism to the balance of trypsin and A1AT in the serum that bathes them constantly? We know that excesses and deficiencies in the levels of either trypsin or A1AT are associated with disease. We know that cellular metabolism can be influenced through signaling involving protease activated membrane GPCR receptors (PAR1-4). In particular, we know of a receptor called PAR2, which is specifically activated by trypsin, expressed by cells at baseline levels, and upregulated through some feedback involving trypsin-activation. We also know that cells at sites of injury or inflammation produce and secrete trypsin, and that this trypsin can act locally upon cells in a variety of ways, all of which have probably not yet been elucidated. Here, we show that sustained exposure to trypsin induces cells to de-differentiate into a stem-like state. We show that if serum is either not added at all, or added and then washed away (after confluency is attained), during cell culture, all cells exposed to exogenously-added trypsin undergo changes in morphology, transcriptome, secretome, and developmental potential, and transition into a state of stemness, in minimal essential medium (MEM). Regardless of their origins, i.e., independent of whether they are derived from primary cultures, cell lines or cancer cell lines, and regardless of the original cell type used, exposure to trypsin (∼10 µM; ∼250 µg/ml) at a concentration 10-fold lower than that used to separate cells during passaging (∼100 μM), over a period of 24-48 hours, causes cells to (1) become rounded, (2) cluster together, (3) get arrested in the G0/G1 stage of the cell cycle, (4) display increased presence of 5-hydroxymethyl cytosine in their nuclei (indicative of reprogramming), (5) display increased levels of activated PAR2 membrane receptor, (6) become capable of very efficient efflux of drug-mimicking dyes, (7) express factors and/or markers known to be associated with induction and/or attainment of stemness, with predominant expression of Sox-2 within cell nuclei; (8) display overall transcriptomic (RNASEQ) profiles characteristic of stemness; (9) secrete stemness-associated factors such as bFGF, and IL-1β, into the medium, in quantities sufficient to support autocrine function (in certain cases); and (10) display increased conversion of pro-MMPs into activated MMPs in the cell’s secretome. Notably, (11) inclusion of differentiating and/or transdifferentiating factors in the environment of such cells causes them to express markers associated with ectodermal, endodermal and mesodermal cell lineages and/or transdifferentiate into specific cell types, e.g., adipocytes or osteocytes.

Most intriguingly of all, (12) the attained stemness appears to be reversible, i.e., withdrawal of trypsin from the medium prior to addition of any differentiating factors restores cells to their original morphology, also over a period of 24-48 hours. Further, (13) a known PAR2 agonist, and a known PAR2 antagonist, respectively, appear to mimic effects of trypsin addition and withdrawal/inhibition. In addition, (14) in experiments with a particular cancer characterized by high levels of stemness (TNBC; triple negative breast cancer), tissues of all TNBC patients express high levels of the PAR2 receptor, as do cells from a known TNBC-derived cell line. We propose that through their effects on PAR levels, and PAR activation status, the balance of trypsin and A1AT levels in organisms normally regulates cellular potential for differentiation, de-differentiation or transdifferentiation, in a local manner, with the default status being that A1AT inhibits trypsin and keeps cells differentiated, whereas sustained trypsin signaling at sites of injury through local production of trypsin helps to place cells into an intermediate state of stemness from which they can either return to their original differentiated state(s), or undergo factor-dependent differentiation, or transdifferentiation, into specific cell types or lineages. It is also possible that reduction in A1AT promotes regeneration. We present a core (RNASEQ-derived) signature for trypsin-induced stemness in human corneal fibroblasts (HCFs) and cells from a retinal pigment epithelial cell line (ARPE-19), noting that there are commonalities as well as differences between them, which suggests that this core signature will be amended with RNASEQ studies of more trypsin-exposed cell types. Our findings offer a possible explanation for the recent unexplained increase in the preference for serum-free protocols used for induction and maintenance of stemness involving iPSCs and mesenchymal stem cells. Also, our studies suggest a new approach to understanding and exploiting how organisms might use stemness, in adults. Trypsin-dominated serine protease induced reprogramming (SPIR) might offer a more natural, and suitably ‘softer’, method of reprogramming of cellular developmental potential for local regenerative requirements in animal tissues.

## INTRODUCTION

Induced pluripotent stem cells (iPSCs) are generally obtained by dedifferentiating adult somatic cells. Presently, the generation of iPSCs from adult cells is entirely dependent upon the deliberately-engineered expression of four protein factors within cells: Oct4, Sox2, Klf4 and cMyc/OSKM (Takahashi and Yamanaka, 2006; Takahashi *et al*., 2007; Park *et al*., 2008; Werig *et al*., 2007; Takahashi and Yamanaka 2016). Sometimes, viruses are used to deliver genes encoding these factors into cells. However, the integration of the viral genome into the cell’s genome carries a significant risk for the reactivation of cancer-related genes (Liu *et al*., 2013). When viruses are not used, of course, the introduced genes do not become integrated into the cell’s genome. However, there is a concomitant reduction in the probability of joint transient and coordinated expression of all four factors, together with a reduction in the probability of such expression continuing across cell generations. These disadvantages act to counter the potential benefits of the non-use of viruses. The efficiency of generation of iPSCs using these methods thus varies between 4.4 % and 0.001 % of the initial cell population, depending upon the cell type involved, and the method used (Zhou *et al*., 2013; Iseki *et al*., 2016; Bar-Nur *et al*., 2014; Buganim *et al*., 2014; Hou *et al*., 2013; Carey *et al*., 2011; Warren *et al*., 2010; Warren *et al*., 2012).

Notwithstanding the highly justified initial enthusiasm associated with the laudable discoveries that led to the development of the above method(s), it does appear today that the associated risks and complexities, as well as the time-consuming nature of the methods, make it necessary for us to search for a more efficient, economical, risk-free and ethically acceptable method of inducing either pluripotency or multi-potency (Brouwer *et al*., 2016; Borgohain *et al*., 2018; Kogut *et al*., 2018; Apostolou and Stadtfeld 2018; Badieyan and Evans; 2019; Shivashankar 2019; Gagliano *et al*., 2019; Haake *et al*., 2019).

The work presented in this paper holds out the possibility of redirecting such a search in a direction away from the deliberate introduction of genes encoding Oct4, Sox2, Klf4 and cMyc/OSKM, towards something that lies potentially upstream of the expression of these four factors, in respect of regulatory mechanisms causing some of the Yamanaka factors to be expressed naturally. Below, we present detailed evidence to demonstrate that the sustained exposure of cells to a serum protein, trypsin, in the absence of any serum, transforms cells into a state of stemness displaying evidence of what could prove to be multipotence, or even pluripotence, with apparent involvement of the PAR2 receptor. The method appears to work upon all cells examined by us, thus far.

It is interesting that this capability of trypsin has not been noted previously, despite the fact that trypsin and EDTA are routinely used by cell biologists to dislodge and transfer (passage) cells between flasks, during cell culture and maintenance. Presumably, the reason is that cell biologists add serum to cells as soon as cells have been dislodged. Serum contains alpha-1 anti-trypsin (A1AT) which inhibits the action of trypsin, such that standardized methods of cell culture provide no scope for sustained exposure of cells to active trypsin, in the absence or presence of serum. We find that when one starves cells of serum, using only minimal essential medium (Dulbecco’s MEM, or DMEM) instead, while ensuring that there is exogenously-added trypsin present in the medium, a number of interesting effects are observed within 24-48 hours, regardless of whether cells are derived from primary cultures, cell lines, or cancer cells lines, and irrespective of the original cell type, across all cells subjected to this treatment. Cells change their morphology, and behavior, becoming spherical and clustering together. There is greater conversion of pro-MMP-2 into MMP-2, and pro-MMP-9 into MMP-9, in the extracellular medium. Cells express known pluripotence-inducing factors, as well as known stem cell markers. Cells display transcriptomic profiles which are similar to those of stem cells, and unlike those of differentiated cells. Cells become efficient at effluxing dyes, similar to stem cells which become chemotherapy-resistant. Cells become arrested in the G0/G1 stage of the cell cycle and display more 5hmc in the nucleus, suggestive of chromatin reorganization. Cells produce factors such as bFGF (basic fibroblast growth factor) and IL-1β, and also secrete these factors into the medium, in quantities similar to those present in standard stem cell maintenance media, and in a manner that depends upon their initial cell type. Cells display the potential for trans-differentiation into other cell types, by the action of specific factors (e.g., into adipocytes, or osteocytes). Cells also display the potential for giving rise to ectodermal, endodermal and mesodermal cells. Further, cells display an increase in expression of the PAR2 receptor, presumably through the operation of a well-known feedback loop in which sustained levels of active trypsin in a cell’s environment increase PAR2 expression, and also cause some increased expression of trypsin by cells. Use of trypsin inhibitors achieves the opposite effects, affecting trypsin expression as well as PAR2 receptor expression.

We are acutely aware of the risks that are associated with the making of claims in this area of work that subsequently prove to be irreproducible. Therefore, we choose to open these findings up to the community at large, for others to confirm, correct, amend or improve, since the necessary experiments are very easy to perform for any group engaged in cellular and molecular biological research. We propose that sustained exposure of cells to active (i.e., non-A1AT-inactivated, or non-aprotinin-inactivated) trypsin gives rise to stemness, and to regeneration of cells and tissues in a manner depending upon the actions of other factors, with serum having the opposite effect of helping to keep cells in the differentiated state(s), at least partially on account of the inactivation of trypsin.

It may be noted that in the interests of maintaining reader interest, in the sections below presenting results and their discussion, we first present experiments that demonstrate the effects of trypsin upon cells, before going on to describe how we arrived at the serendipitous discovery of these effects that have escaped the attention of generations of investigators. We call our new method of generating stemness, through trypsin-induction, SPIR (serine protease induced reprogramming), since there are multiple trypsin-analogs in the serum, as well as proteins like elastase which could all eventually prove to be involved in mediating such effects.

## MATERIALS AND METHODS

### Cells

Cells used include (a) cells in primary cultures of HCFs, i.e., human corneal fibroblasts sourced from corneo-scleral rims discarded during routine keratoplasty for corneal transplants, (b) cell lines, including ARPE-19 (Retinal pigment epithelial cells; ATCC); MCF7 and MDA-MB-231 (breast cancer cells), and NCI-H292 (lung cancer cells), and (c) cells in tissues derived from surgery of triple-negative breast cancer patients.

### Culture conditions

Initially, cells were maintained in DMEM medium with 10 % FBS at 37 °C and in 5 % CO2. Subsequently, with the temperature and CO2 conditions remaining the same, cells were cultured under one of the following types of conditions, after they had reached a state of 80 % confluency: (i) SS (serum-starved) conditions, where serum-containing medium was replaced with serum-free DMEM, in routine cell culture; (ii) SF (serum- and growth factor-free) conditions, where no serum was used after cells were subjected to trypsinization during passaging, and instead cells from multiple cultures were pooled, washed once with PBS, and then plated out at high density (∼1×10^6^ cells/ml) in DMEM for 24-96 h; (iii) SFA (SF with aprotinin) conditions, where the bovine pancreatic trypsin inhibitor (also known as BPTI, or aprotinin) was added to cells being cultured under SF conditions, and (iv) SPIR (serine protease-induced reprogramming) conditions, where serum-containing medium was replaced with serum-free DMEM for 24 h, as in the SS condition, and this was followed by the exogenous addition of trypsin, for 24-48 h, by exposing cells to a ten-fold dilution of the same stock of trypsin originally used for passaging (addition of 1 ml of trypsin-EDTA stock to 9 ml of DMEM) bringing down the concentration of trypsin from that used for passaging, from 0.25 % w/v (2.5 mg/ml; 100 μM) to 0.025 % w/v (0.25 mg/ml; 10 μM). Variations of SPIR included SPIR-W which included withdrawal of trypsin following SPIR, and SPIR-F which included addition of differentiating factors to cells being subjected to SPIR.

### Zymography

Gelatin (1 mg/ml) was co-polymerized in the lower separating gel of a 10 % SDS-PAGE gel. Samples for electrophoresis were prepared with non-reducing sample loading buffer (NRSLB). SDS was removed from the gel by incubation with 2.5 % Triton X-100 for 1 h, followed by overnight incubation in activity buffer at 37 °C. The gel was stained with Coomassie Brilliant Blue G-250 and then de-stained, to visualize the zone of clearance arising from any gelatinolytic activity.

### Fluorogenic assay(s)

Matrix metalloprotease (MMP) activity in test samples was assessed by quantitating the fluorescence of Mca-KPLGL-Dpa-AR-NH2, a fluorogenic peptide substrate of MMPs (R&D systems, USA). The fluorgenic substrate (5 μM) was mixed with control and test samples in 96 well plates (Black wall plate, Nunc), incubated for 1 h at 37 °C, and examined for fluorescent product (excitation/emission: 320/405 nm).

### Immunofluorescence studies

Cells were grown in a 24 well plate with sterile glass coverslips. After SPIR treatment, care was taken not to abstract treated cells, as cells aggregated into spheres loosely bound to coverslips. Cells were washed and fixed with 4 % paraformaldehyde solution for 10 min, blocked and permeabilized using 0.3 % Triton X-100, 1 % BSA and 0.3 M Glycine, respectively, at room temperature. Primary antibodies against proteins of human origin including Oct 3/4, Nanog, SOX-2, 5hMC, Tra-1, Ki67, and the cell surface protein, PAR2, were used. Cells were incubated overnight with a primary antibody in a refrigerator, at 2-8 °C. On the next day, cells were washed with PBST (PBS containing 0.1 % BSA and 0.2 % Tween 20). Subsequently, cells were incubated with a suitable (fluorochrome labelled) secondary antibody for 45 min at room temperature. After washing with PBST, actin labelling was done with TRITC-labelled Phalloidin, followed by counterstaining with DAPI, and mounting was done with 90 % glycerol on a clean glass slide. Image acquisition was done using an objective lens of 63x magnification and 1.4 numerical aperture on a confocal laser scanning microscope (TCS SP8, Leica Microsystems, Germany). No primary antibody controls were used to check any nonspecific staining and all the images were acquired in sequential imaging mode to remove any spectral bleeding effects. All immunofluorescence experiments were repeated at least three times on separately cultured specimens.

### Cytokine bead array assay

H-CBA (human cytometric bead array) assay was performed for quantification of various cytokines and growth factors, using the protocol provided by the manufacturer. Serial dilutions of human flex cytokine/growth factor standards were prepared using assay diluents. Prior to use, capture beads were briefly vortexed. Then, 50 µl of the bead suspension was added to all labelled assay tubes, followed by addition of 50 µl of human standards to the control assay tubes, 50 µl of each test sample to the appropriately labelled sample assay tubes, and 50 µl of PE detection reagent to all assay tubes, followed by incubation for 2 hours at room temperature. Assay tubes were washed with wash buffer and centrifuged at 200 *g* for 5 minutes; the wash buffer was carefully aspirated. Beads were re-suspended in 400 µL of wash buffer and measurements were made using BD FACS ARIA.

### Cell cycle stage analysis

Cell cycle stage analysis was done using flow cytometric estimation of cellular DNA bound to the dye, propidium iodide (PI). Cells were harvested using trypsin-EDTA solution and centrifuged at 250 *g* for 5 minutes. After two washes with PBS, the cell pellet was dispersed using the finger-tapping method, followed by addition of ice-chilled 70% ethanol. Cells were stored in the refrigerator prior to flow cytometric analysis on a BD Accuri C6 instrument (Becton-Dickinson, USA). Before flow cytometry, ethanol was removed and cells were washed using PBS and resuspended in PBS containing RNase (50 µg/mL) and PI (50 µg/mL). Doublets were removed using the Area vs. Height plots for PI parameter. Cell cycle distribution was represented as a histogram of DNA:PI.

### Drug efflux assay

Rhodamine 123 was used for measurement of total cellular efflux activity. Cells were harvested using trypsin-EDTA solution, washed twice with PBS and centrifuged. The cell pellet was resuspended in 1 ml of ice-cold DMEM medium containing 1 % bovine BSA, spiked with Rhodamine 123 in DMSO. Cells were incubated for 30 min on ice with the dye. After incubation, cells were washed with DMEM to remove free dye, and the rhodamine-loaded cells were aliquoted and kept either on ice or in a thermostatically regulated water bath (37 °C) for 1 h. Cells were then washed twice, resuspended in DMEM medium and immediately analyzed on a BD Accuri C6 flow cytometer (Becton-Dickinson, USA). All samples were kept on ice, and were removed from ice just prior to data acquisition on the flow cytometer.

### Transcriptome profiling using RNA quantification sequencing

RNA was isolated from cells (ARPE19, or HCF) under SPIR and SS conditions, and processed for next-generation sequencing (NGS) to generate deep coverage RNASeq data. For each treatment group, sequencing was performed on at least 2 biological replicates. Sequencing libraries of Poly-A enriched RNA were generated using the NEBNext® Ultra™ II RNA Library Prep kit, according to the manufacturer’s protocol. Library quality control was assessed using the Agilent DNA High Sensitivity Chip and qRT-PCR. High quality libraries were sequenced on an Illumina NextSeq 500 instrument. To achieve comprehensive coverage for each sample, we generated ∼25-30 million paired-end reads.

### RNASEQ data analysis

The raw sequencing data was processed to remove any adaptor, PCR primers and low quality transcripts using FASTQC and Trimomatic, which provide a very comprehensive estimate of sample quality on the basis of read quality, read length, GC content, uncalled bases, ratio of bases called, sequence duplication, adaptor and PCR primer contamination. These high quality, clean reads were further aligned against human genome using tophat2 and bowtie2 packages (http://tophat.cbcb.umd.edu/). The Grch37 human genome assembly was used as a reference genome for the alignment. Gene expression measurements were performed from aligned reads by counting the unique reads. The read count-based gene expression data was normalized on the basis of library complexity and gene variation using the R package Voom. The normalized count data was compared among groups using a linear model to identify differentially expressed transcripts. The differentially expressed transcripts were identified on the basis of assessment of multiple test-corrected p value and fold change. Genes were considered significantly differentially expressed if the multiple test corrected p-value was < 0.01 and the absolute fold-change was > 2.0. The SPIR core transcriptome signature was generated from genes that are consistently up- or down-regulated by SPIR treatment in both ARPE19 and HCF cell types.

### Functions, pathways and interactive network analysis

Ingenuity Pathway Analysis (IPA 9.0, Qiagen) was used to identify the pathways that are significantly affected by transcripts that are significantly differentially expressed due to SPIR treatment in different cell types. The knowledge base of this software consists of functions, pathways and network models derived by systematically exploring the peer reviewed scientific literature. A detailed description of IPA analysis is available at the Ingenuity Systems’ web site (http//www.ingenuity.com). The approach calculates a p-value for each pathway according to the fit of users’ data to the IPA database, using the one-tailed Fisher’s exact test. The pathways with multiple test corrected p-values < 0.05 were considered significantly affected. Further the tool also predicts activation or inhibition of pathways by calculate a z-score based on match between expected relationship direction and observed gene expression changes.

### Regulator Module Analysis

Regulatory module analysis was used to identify the cascade of upstream transcriptional regulators involved in the gene expression changes observed, to help identify key regulators (master regulators) and understand the underlying biological mechanisms. The analysis helps in identifying the transcriptional regulators that are significantly affected by the SPIR treatment. It also helps in determining whether the regulators are activated or inhibited based on Z-score calculations. The significance of overlap was determined using the one-tailed Fisher’s exact test.

### Gene Set Enrichment Analysis (GSEA)

GSEA was performed using the GSEA-R, a Bioconductor implementation of GSEA from the Broad Institute, Massachusetts, USA. The GSEA analysis was performed to compare the transcriptome signatures of SPIR treated cells various stem cell signatures in MsigDB 6.0. The enrichment transcriptome signature of SPIR in stems cell gene-sets was performed by developing a customized dataset of gene-sets related to stem cells from MsigDB 6.0. The analysis was performed with 1000 permutations and a classic statistic; NES = normalized enrichment score and Nominal P value was measured.

### Comparative analysis with external stem cell dataset

To determine the enrichment of the stem cell phenotype in SPIR treated samples, we performed enrichment analysis of SPIR signatures of each cell into an external dataset consisting of information concerning the transcriptomes of differentiated cells and stems cells. The SPIR core transcriptome signature was generated from genes that are consistently up- or down-regulated by SPIR treatment in both ARPE19 and HCF cell types. Transcriptome dataset of comparing stem and differentiated cells were obtained from the GEO database (i.e. GSE12390) and normalized before performing the GSEA analysis. Enrichment analysis was run with 1000 permutations and a classic statistic; NES = normalized enrichment score and Nominal P value was measured. Further we also performed enrichment analysis of SPIR core signature in collection of stemness signature and datasets available in StemChecker tool [PMID 26007653].

### F2RL1 association with Survival across major TCGA cancer types

To determine the association of key SPIR altered genes with survival in various cancers, we performed survival analysis using RNASEQ data from ∼10000 patients in the TCGA database (PMID: 24071849), by looking at the pattern of individual gene expression in each cancer. The expression data was divided into high, medium and low expression groups on the basis of expression profiles. We performed survival analysis using the multivariate Cox proportional hazards model and log-rank tests based on Kaplan-Meier curves. The survival analysis was carried out using R survival package, which estimates the hazard ratios (HR) for each gene, after adjustment for age, gender and cancer stage, when such information is available. The results were considered significant if the log rank test p-values were < 0.05. Additionally, to explore association of F2RL1 gene with aggressiveness in the breast cancer, we also evaluated its expression profile in triple negative (TN) vs. non-triple negative (Non-TN) breast cancer samples in an external microarray dataset (i.e. GSE76275). The significance of difference in expression profile between TN vs. Non-TN was determined using t-test in R.

### Patient selection and biopsy

This was a retrospective study which included 60 patients who had undergone mastectomy/lumpectomy with axillary clearance for carcinoma breast. All cases were evaluated for ER, PR and Her2 expression and cases with triple negativity were enrolled in the study. The patients were divided into 2 groups of 30 patients each; one group of 30 patients with no lymph node involvement and the other with 30 patients with lymph node involvement. Ethical clearance from the Institutional Ethics Committee of the Postgraduate Institute of Medical Education and Research (PGIMER, Chandigarh) was obtained.

### Immunohistochemical staining

Tissue sections of 4 to 5 μm thickness were mounted on freshly prepared 0.01 % poly-L-lysine coated slides, and fixed overnight at 37 °C. The slides were dewaxed and rehydrated in a graded series of alcohol solutions. Endogenous peroxidase activity was blocked by immersing coated sections in freshly prepared 3 % hydrogen peroxide in methanol for 20 min, followed by three washes in PBS. Antigen retrieval was done by the heat retrieval method, using the automated DAKO antigen retrieval system. The primary antibodies against ER, PR, HER2 and Ki67 were diluted according to standardized protocols and applied to tissue sections. PAR2 antibody diluted in BSA was applied to the sections in a dilution of 1:50 and kept at room temperature for 90 minutes in a moist chamber. Incubation with the secondary antibody, Dako Envision, was done for ER, PR, HER2 and Ki67 according to standardized protocols. The sections were covered with freshly prepared diamino benzidine, or DAB (1 ml of substrate, hydrogen peroxide with one drop of the chromogen, DAB) followed by counter staining with haematoxylin. The sections were then dehydrated, cleaned in xylene and mounted with DPX.

### Karyotyping analysis of SPIR-treated ARPE-19 cells

Karyotyping analysis was performed to detect any chromosomal aberrations. Actively dividing ARPE19 cells, under SS and SPIR treatment conditions, were exposed to 0.1 µg/mL colcemid solution at 37 °C. After 2 h of colcemid exposure, cells were detached using 0.25 % trypsin, 1mM EDTA, solution and centrifuged for 10 min at 250 xg. Both cell pellets were dispersed by light tapping and hypotonic treatment was given by adding 10 mL of 0.075 M potassium chloride for 12 min at 37 °C. Post hypotonic treatment, cells were centrifuged at 200 xg for 5 min. The cell pellets were dispersed and fixed by the addition of ice-chilled Carnoy’s fixative (Methanol: Acetic Acid 3:1). Clean glass slides were used for slide preparation by the drop-fall method and immediately placed on a hot plate for 2 h at 45 °C. Heated slides were dipped in ice-cold 0.37 % trypsin solution for 8 s and immediately washed twice with cold water. Giemsa staining in 10 % v/v Giemsa stain was performed for 3.5 min. Stained slides were visualized on an upright microscope at 100 X magnification, and images were captured using a CCD camera (Olympus, Japan). At least 10 metaphases were analyzed using GenASIs Bandview software (Applied Spectral Imaging, USA).

## RESULTS AND DISCUSSION

We begin this section with a short re-introduction of different cell culture conditions used in this work, since we feel that a clear understanding of these is essential for comprehension of the data presented below. The 1^st^ culture condition is called SS (serum starved). In SS, cells are dislodged through the addition of trypsin-EDTA after they have reached 80 % confluency, then exposed to serum to inactivate the trypsin, and then transferred to a new flask, and allowed to reach 80% confluency again in the presence of serum, and further washed with PBS to remove serum, and thereafter maintained in Dulbecco’s minimal essential medium (DMEM) and never again exposed to either serum or exogenously-added trypsin. The 2^nd^ culture condition is called SF (serum- and growth factor-free). In SF, we transfer trypsin-EDTA treated (and dislodged) cells into a new flask without the addition of any serum to inactive the trypsin at any stage. Thereafter, we maintain the cells in DMEM. Notably, in SF, we also pool cells from many initial trypsin-EDTA treated cultures in order to plate them out at high densities in DMEM, thus ensuring scope for the presence (and activity) of residual trypsin from the trypsin-EDTA treatment. The 3^rd^ culture condition is called SFA (SF with aprotinin added). In SFA, conditions are identical to SF, except that aprotinin (pancreatic trypsin inhibitor) is added. The 4^th^ culture condition is called SPIR (serine protease induced reprograming). In SPIR, the same protocol used in SS is again used, except that addition of trypsin to the DMEM follows serum starvation, as detailed in the methods section and as also shown in a schematic diagram (Fig 1). Notably, the trypsin used for sustained exposure in SPIR is a 10-fold dilution of the same stock of trypsin-EDTA used for passaging of cells, used over 24-48 h instead of over 5-10 minutes. The 5^th^ culture condition is called SPIR-W, in which trypsin is withdrawn/removed after SPIR, through changes of DMEM. The 6^th^ culture condition is called SPIR-F in which SPIR is used with transdifferentiating factors, in experiments intended to differentiate cells that have undergone de-differentiation through trypsin exposure.

**Fig 1:**
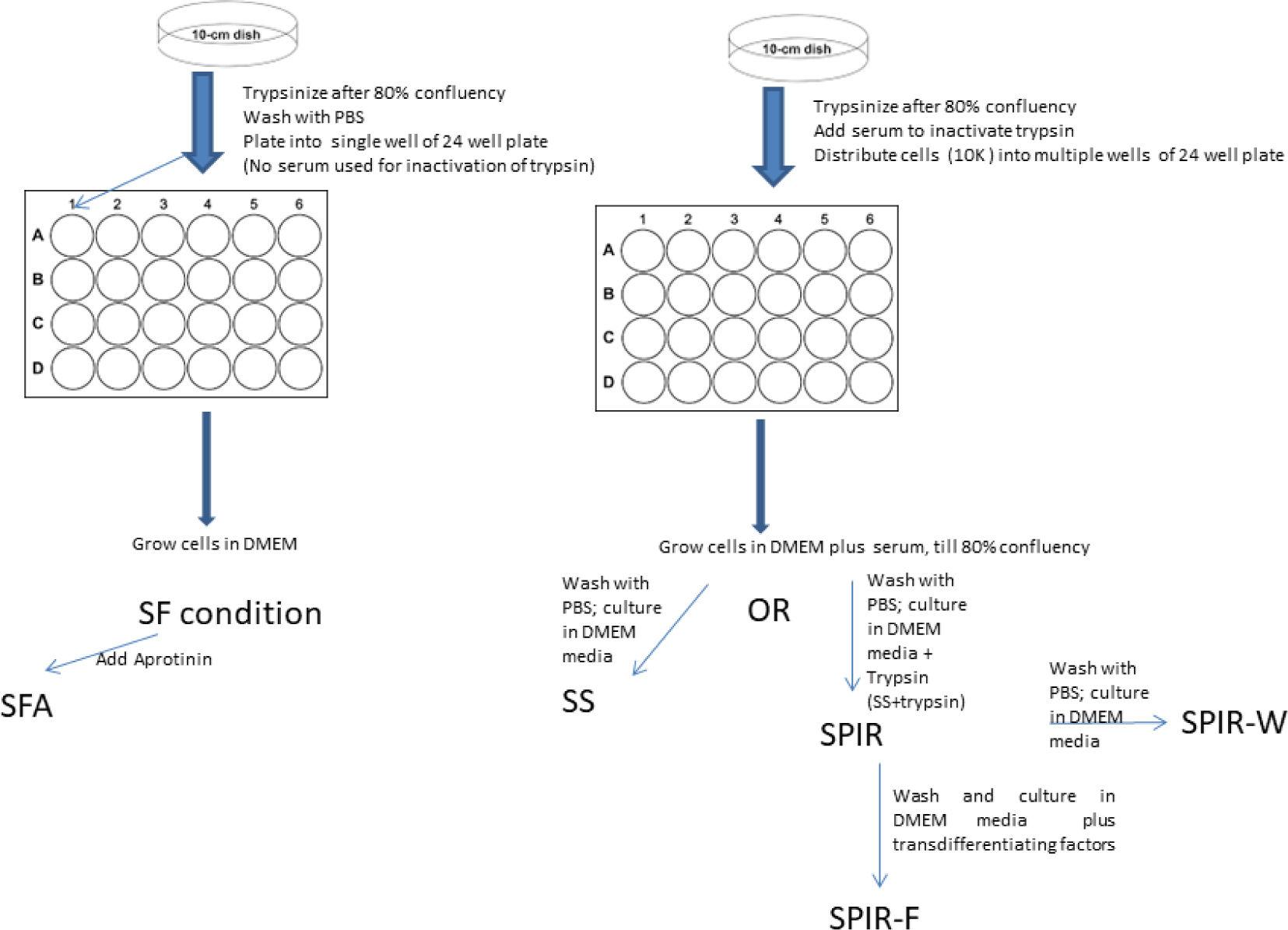
Schematic diagram showing the various methods used to culture cells (detailed in the “Results and Discussion” section).

### Cytokine Bead Array (CBA) profiling of the secretome under SS and SF conditions

In an attempt to characterize cells cultured under SF conditions, the human cytometric bead array (H-CBA) assay was carried out, using conditioned medium (CM) from cells cultured under SS and SF conditions. The secretomes of ARPE-19, MDA-MB231, NCI-H292 and HCF cells were analyzed for the presence of various cytokines and growth factors. As seen in Table 1, ARPE-19 and HCF cells showed increased levels of both bFGF and IL-1β under SF conditions.

**Table 1:** Secretome analysis of conditioned media (25 X concentrated) from ARPE19 cells cultured under SF and SS conditions by cytokine bead array

| Analyte | SF (pg/ml) | SS (pg/ml) |
| --- | --- | --- |
| Human IL-8 | 2891.3 ± 2119 | 5071 |
| Human IL-1β | 372.0 ± 283.8 | NA |
| Human TGF-β1 | 306.9 ± 250.9 | 618.6 |
| Human VEGF | 1479.9 ± 390.6 | 1904.6 |
| Human MCP-1 | 9061 ± 3806 | 14649.2 |
| Human IL-12 | 19.3 ± 28.1 | 4.7 |
| Human bFGF | 250000.0 ± 56.6 | NA |

### Immunofluorescence microscopic profiling of the secretome under SS and SPIR conditions

In ARPE-19 cells cultured under SF conditions, co-staining of bFGF and lysotracker dye (a fluorescent weak base that accumulates in highly acidic organelles like the lysosome) suggests that bFGF and IL-1β are produced and localized in acidic vesicles (Supplementary Fig 1). MDA-MB231 and NCI-H292 cells did not show expression of bFGF; however, IL-1β was seen to be significantly increased in MDA-MB231 cells (data not shown). The finding of the upregulation of these two proteins was intriguing, as both bFGF and IL1-β tend to be associated with upregulation of stem cell gene expression (Wang *et al*., 2005; Xu *et al*., 2005; Lanner and Rossant, 2010; Greber *et al*., 2010; Brady *et al*., 2013). The SPIR condition described was specifically developed to facilitate the exploration of trypsin involvement, in comparison with the SS condition, in a systematic manner applicable to all cells, in this and subsequent experiments. Noting the increased expression of soluble bFGF and IL1-β, we decided to further explore the importance of trypsin and any connection that it has to stemness, in greater detail, beginning with RNASEQ analyses.

### RNASEQ profiling of cells cultured under SS and SPIR conditions

We decided to carry out RNA sequencing (RNASEQ) analyses of two different cell types (ARPE-19, a non-cancerous cell line, and HCF, a primary culture of fibroblasts) under SS and SPIR culture conditions, mainly to examine through transcriptome analyses whether trypsin confers stemness. The transcriptome profiling shows that SPIR culture conditions significantly altered 1,614 and 2,125 genes in ARPE-19 and HCF cells, respectively (Absolute Fold Change ≥ 2; multiple test corrected P value < 0.05). The heatmaps of top SPIR-altered genes in ARPE-19 and HCF cells are shown in Fig 2A and Fig 2B respectively. Further comparative evaluation of gene expression in ARPE-19 cells, and HCF cells, upon SPIR treatment, showed that the common expression signatures under SPIR culture conditions (which we shall call the SPIR core signatures) consistently displayed upregulation of 337 genes and downregulation of 380 genes in both cell types, in relation to expression under SS conditions (Fig 2C).

**Fig 2:**
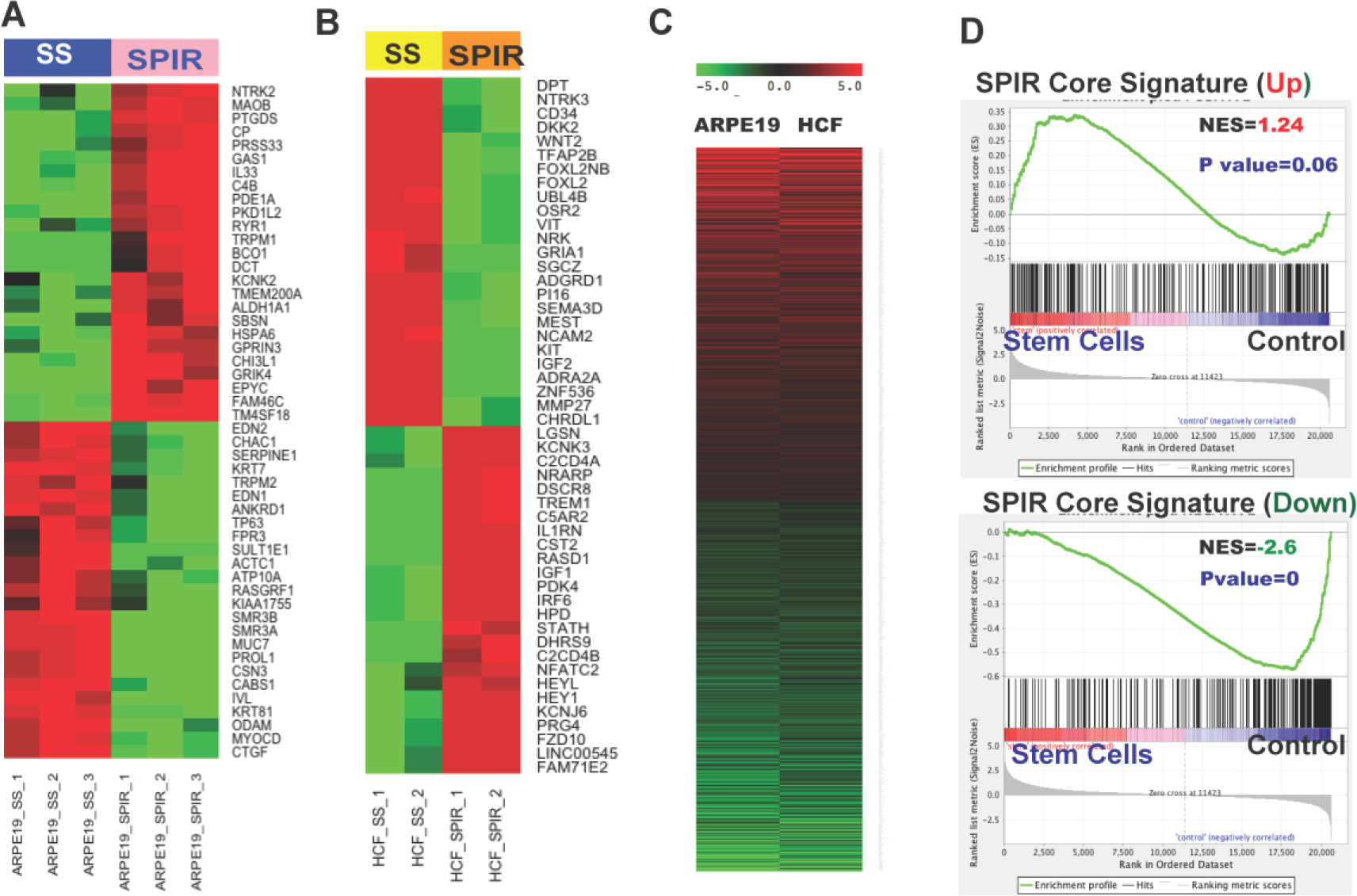
Transcriptome profiles of ARPE-19 cells and HCF cells upon SS and SPIR treatments. Heatmaps of top genes that are differentially expressed in (A) ARPE19 cells, and (B) HCF cells, under SPIR culture conditions relative to SS culture conditions, are shown. In the heatmaps, rows depict differentially expressed genes and columns depict samples cultured in SPIR and SS conditions. (C) Fold-change based heatmap of the core set of genes which are modulated consistently in ARPE-19 and HCF cells by SPIR culture conditions. Red and green colors represent upregulated and downregulated genes, respectively. (D) Positive enrichment analyses of SPIR-upregulated core signature(s) with external datasets comparing stem cells with control/differentiated cells (upper panel). Negative enrichment analysis of SPIR-downregulated core signature(s) with external datasets comparing stem cells with control/differentiated cells (bottom panel).

Enrichment analyses of the SPIR core signatures were performed against an external dataset (GEO database: GSE 12390) which consists of transcriptome information from stem cells and differentiated cells. The analyses yielded strong enrichment scores of 1.249 and −2.54, with p values corresponding to 0.06 and 0.0, respectively, for the upregulated and downregulated genes, respectively, for stem cells (Fig 2D). These results indicate that the SPIR culture conditions upregulate the genes linked with stemness and downregulate the genes linked with cellular differentiation.

The pathways analyses on genes that are significantly altered due to SPIR culture conditions show activation (based on positive Z-score) of the acute phase response signaling and glycolysis in ARPE-19 cells, and activation of TGF-β, Notch, IL-6, p38, MAPK and the NRF2-mediated oxidative stress response pathways, in HCF cells (Fig 3A). To gain further insight into the upstream key regulators that might be responsible for observed changes after SPIR treatment, we performed a systems biology-oriented analysis. The regulator analysis depicted significant activation of VEGF, NOTCH, MAP2K1, CREB in HCFs (Fig 3B). The significance of such pathways has been discussed in a later section.

**Fig 3:**
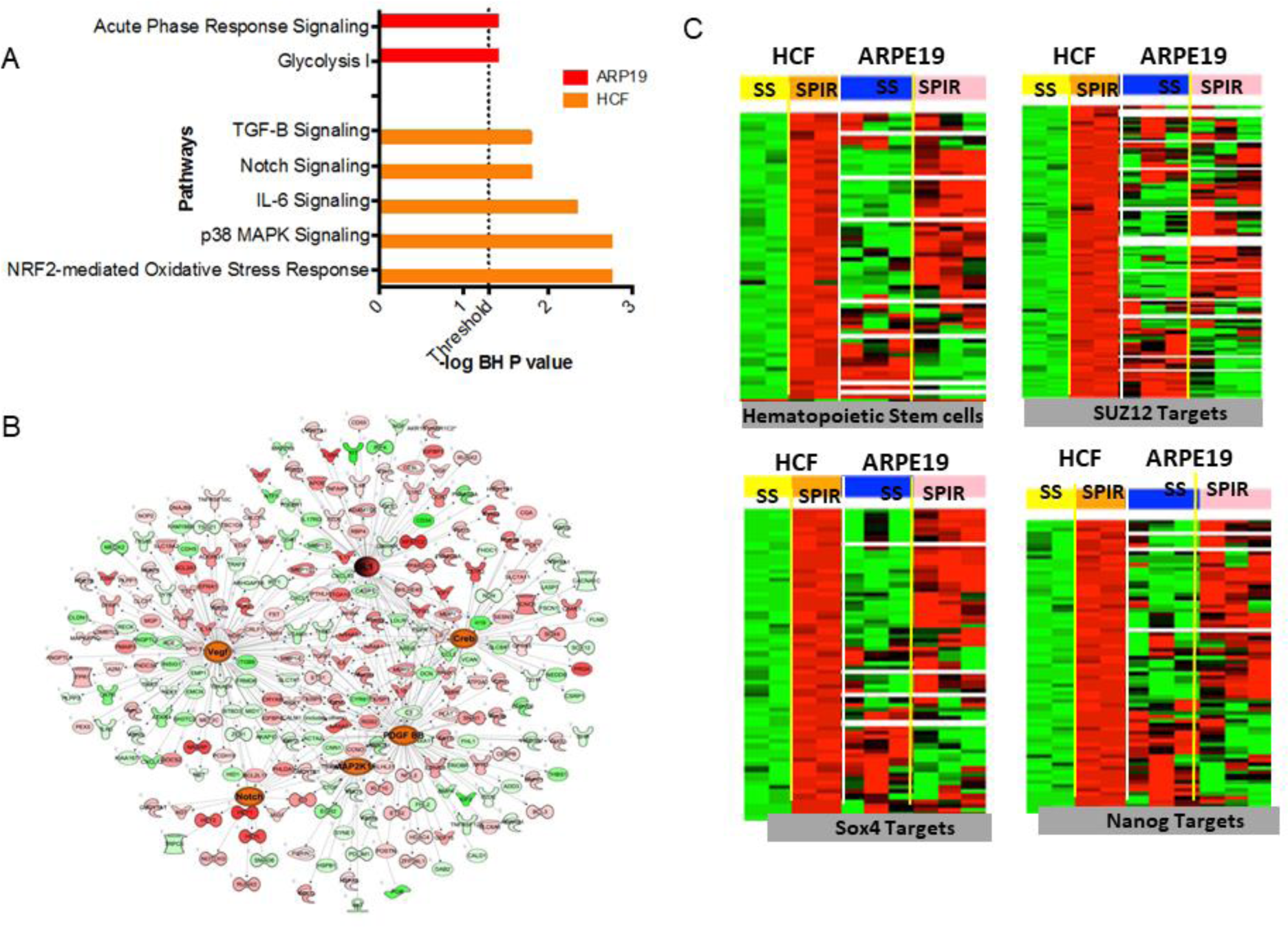
(A) Enrichment map of pathways that are significantly activated by SPIR treatment in HCF or ARPE19 cells. The Y– axis represent significantly effected canonical pathways and X-axis-log transformed P value. All these pathways are predicted activated based on Z score >1.5. (B) Network of top activated regulators and their target genes in HCF. (C) Heat maps for enrichment of SPIR upregulated genes in HCF and ARPE19 analyzed with respect to stemness signatures and key transcription factors related to stemness.

Further stemness-focused related-enrichment analyses were performed for both cell types against stem cell signatures, as well as in regard of key transcription factors related to stemness. It was observed that whereas SPIR conditions caused HCF cells to express genes related to embryonic stem cells, neural stem cells, hematopoietic stem cells and mammary stem cells, the p value was most significant for the latter two stem cell types (Fig 3C).

In the case of ARPE-19 cells, SPIR culture conditions caused cells to express genes most closely related to embryonic stem cells and neural stem cells (Table 2). There was also a significant increase in levels of mRNA encoding key transcription factors known to be related to stemness (Table 3); the heat map for the analysis, as already mentioned, is shown in Fig 3C.

**Table 2:**
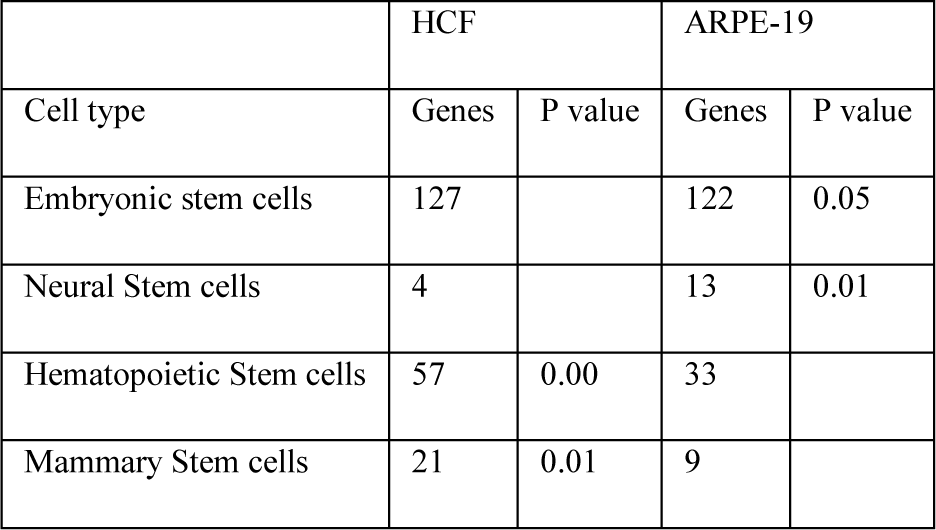
Enrichment of SPIR modulated genes in the canonical stemness signatures.

**Table 3:** Enrichment of SPIR modulated genes in the key stemness related transcription factors.

| TF Name | HCF | ARPE-19 |
| --- | --- | --- |
|  | P value | P value |
| Nanog | 0.014 | 0.01 |
| SOX-2 | 6.76E-04 | 0.013 |
| OCT-4 |  | 0.065 |
| SUZ12 | 3.01E-10 | 4.60E-13 |
| SMAD4 | 0.001 | 0.005 |
| SMAD2 | 0.002 | 4.18E-04 |
| SMAD3 |  | 0.001 |

In the following sections, we describe experiments characterizing stemness induced by SPIR culture conditions using other approaches.

### Morphology of cells under SPIR and SPIR-W conditions

Under SPIR culture conditions, cells displayed a tendency to change into a round/spherical shape, regardless of the original shape and morphology. Cells were also found to display a tendency to cluster, or clump together, over a time span of a few hours to a few tens of hours (typically 24-48 hours). Interestingly, when SPIR conditions were withdrawn, i.e., when trypsin was withdrawn/removed from the culture during changes of the DMEM (SPIR-W), cells were once again found to display their characteristic original differentiated morphologies after a further 24-48 hours, following the withdrawal of trypsin (Supplementary Fig 2-4). Given that SPIR-treated cells appear to have transitioned into a state of stemness, clearly the ability to return to the original morphology suggests that this is a different kind of stemness from anything seen previously. We performed karyotyping analysis (data not shown) to satisfy ourselves that SPIR treatment does not result in any chromosome breaks, or rearrangements in chromosome banding patterns, using ARPE-19 cells.

Stem cells do not characteristically differentiate back to their original cell type, e.g., if they are induced pluripotent stem cells. Nor do totipotent cells created through nuclear transplantation into fertilized eggs differentiate back from their undifferentiated state into the original cell type from which the nucleus was derived, e.g., a skin cell. The level of reprogramming that occurs after nuclear transplantation and growth of a zygote would appear to be absolute, in the case of totipotent stem cells. It is also near-absolute in the case of induced pluripotent stem cells and other kinds of stem cells, with little scope present for the formation of only one cell type through further growth and division of the stem cells, without the cells going through multiple cell lineages.

On the other hand, here it is observed that when trypsin is withdrawn from the medium, and no differentiating factors are added during the period of exposure of cells to trypsin (i.e., during SPIR treatment), there is apparently only some limited nuclear and cellular reprogramming occurring. This is because a return to the original morphology would otherwise have been impossible, were cells to have been reprogrammed to a more profound level, erasing memory of their immediate past. Therefore, it appears that trypsin causes cells to be only partially-reprogrammed into a state of stemness; one in which cells do not forget their origins, but are yet clearly dedifferentiated as well as amenable to redifferentiation and transdifferentiation. We emphasize, therefore, that the stemness observed here is a different state of stemness from that of iPSCs, and that it constitutes a new state of stemness which could be subtly different for different cell types, with some common features. We also emphasize that many investigators working with different cell types in cell culture would have probably observed that cells become rounded and tend to cluster and survive, after prolonged exposure to trypsin in the absence of serum. They would have also observed that such cells manage to return to their original differentiated states upon the addition of serum, without ever realizing that the cells had begun the process leading to attainment of a certain state of intermediate stemness, but happened to be pulled back through serum addition.

### Immunofluorescence confirmation of stem cell marker expression during SPIR

Sox-2, Nanog and Oct4 are known stem cell markers. Immunochemical staining for expression of these stem cell markers was strongly positive, under SPIR culture conditions, in multiple cell types (Supplementary Fig 5, 6). In MCF7, ARPE-19, HCF and NCI-H292 cells, i.e., experiments covering cancer cell lines (MCF7 and NCI-H292), a non-cancerous cell line (ARPE-19) and primary cultures (HCF; human corneal fibroblasts), similar changes were seen in respect of upregulation of Sox-2, Oct-4 and Nanog.

**Fig 4:**
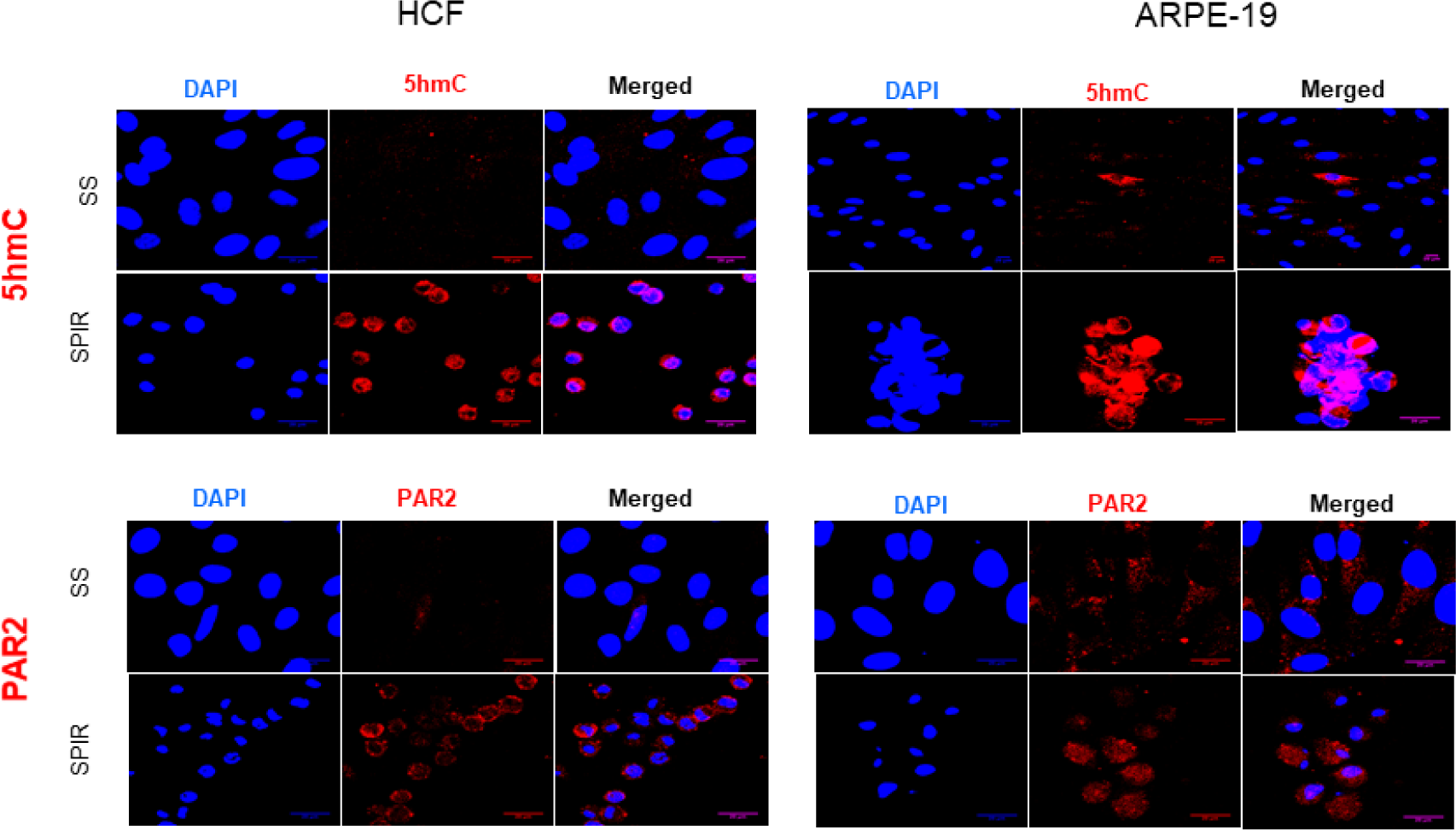
Immunostaining of ARPE-19 cells and HCFs for localization of 5hmC and PAR2 receptor expression. DAPI nuclear staining (blue), phycoerythrin (PE) - labeled 5hmC or PAR2 (red), and merged fluorescent images were captured by confocal microscopy (scale bar = 20 μm).

**Fig 5:**
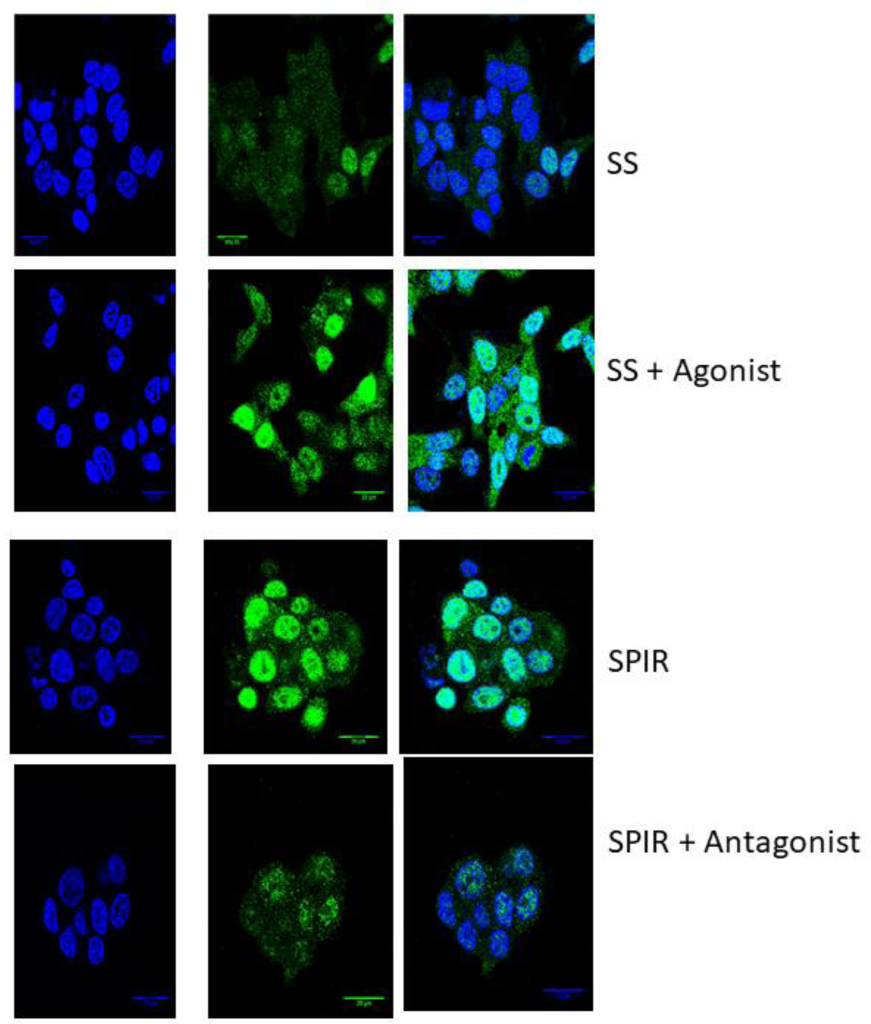
Sox-2 expression in MCF-7 cells seen under SS conditions, SS conditions supplemented with PAR2 specific agonist, SPIR conditions and SPIR conditions supplemented with PAR2 antagonist. DAPI nuclear staining (blue), FITC-labeled Sox-2 (green), and merged fluorescent images were captured by confocal microscopy (scale bar = 20 μm).

### Increased PAR2 receptor presence during SPIR

To examine whether trypsin has an effect upon the cell surface receptor, protease activated receptor 2 (PAR2), we observed the levels of expression of PAR2 in cells cultured under SS and SPIR conditions. Fig 4 shows that expression of the PAR2 receptor is significantly increased under SPIR culture conditions in HCF and ARPE-19. Supplementary Fig 7 shows that expression of the PAR2 receptor is significantly increased in NCI-H292 and MCF7 cells.

### Increase in nuclear 5-hydroxymethylcytosine (5hmc) during SPIR

We also investigated the levels of 5hmC in the nuclei of different types of cells (ARPE-19, MCF-7, HCF and NCI-H292) cultured under SS and SPIR conditions. Clearly, 5hmc could be observed to be highly enriched in the nuclei of SPIR-treated cells (Fig 4; Supplementary Fig 7), in comparison to cells cultured under SS conditions. Increased 5hmc in the nucleus is suggestive of alterations in the global

DNA demethylation status and chromatin reorganization (Shi *et al*., 2017; Kumar *et al*., 2018). This suggests that SPIR stimulates nuclear reprogramming, which we have already speculated above is likely to be a more limited reprogramming than that applicable to iPSCs, because of the reversibility of such reprogramming. In all likelihood, SPIR-treated cells are like some of the intermediate states seen during progression of cells into iPSCs.

### Mimicry of SPIR and SS conditions by PAR2 agonist/antagonist

We also used a known agonist (peptide Ser-Leu-Ile-Gly-Lys-Val-NH2, **SLIGKV-NH2**) as well as a known antagonist of PAR2 (ENMD 1068, Santa Cruz), in attempts to mimic SPIR and SS culture conditions, since an agonist could mimic the effect of trypsin, and an antagonist would block trypsin-induced signaling through PAR2. We examined expression of a representative stem cell marker, Sox-2, with the agonist and the antagonist (Fig 5). Use of the agonist under SS culture conditions (attempting to mimic SPIR culture conditions) results in increased expression of Sox-2 in MCF7 cells. Notably, thrombin which is recognized by the PAR1 and PAR4 receptors does not elicit any increase in Sox-2 expression (data not shown). Use of the antagonist, however, decreased the basal level of Sox-2 expression in cells cultured under SPIR conditions. Thus, both when trypsin or a PAR2 agonist are used, there is increase in the expression of PAR2. It is known that PAR2 expression increases when there is trypsin present in the environment, ostensibly on account of a feedback mechanism regulating PAR2 expression and activation through availability of its ligand (Darmoul *et al*., 2001).

### Reversible arrest of cells in the G0/G1 stage of the cell cycle during SPIR

To assess the effect of SPIR on cell cycle and proliferation, we analyzed cell cycle status using propidium iodide-based flow cytometry, after culture under SS and SPIR conditions. We also analyzed cell cycle status after withdrawal of SPIR conditions (SPIR-W). We observed using flow cytometry that SPIR treatment causes all cells to become arrested in the G0/G1 phase of the cell cycle, in both MCF7 and ARPE cells, whereas under SPIR-W conditions, cells once again distribute out into populations in the S, G2, and M phases of the cell cycle (Supplementary Fig 8A). Further, cells were labelled with the Ki67 antibody that is cognate to the nuclear protein, MK167, a ribosomal RNA transcription factor which is also a cell proliferation antigen and marker. It was observed that the expression of MK167 is downregulated under SPIR conditions in comparison with both SS culture conditions, and SPIR-W conditions (Supplementary Fig 8B). The nuclear localization of MK167 in SPIR was confirmed using confocal imaging of Ki67-stained cells grown under SS and SPIR conditions (Supplementary Fig 8C).

### Reversible enhancement of drug-efflux during SPIR

It is well established that epigenetic alterations (including DNA methylation and histone modifications) tend to be associated with increased resistance of cells to chemotherapeutic agents, and that this is due to upregulation of the activities of transporters engaged in drug efflux. Therefore, we next studied the activity of a well-characterized transporter protein, ABCB1 (MDR-1/P-gp) in cells subjected to SPIR culture conditions. Using flow cytometric evaluation of efflux of the fluorescent dye rhodamine 123 (used as a reference P-gp substrate probe), we characterized the extent of P-gp activity in cells cultured under SS and SPIR conditions, by measuring the intracellular accumulation of rhodamine 123. In cells of the MCF7 cancer cell line, under SS conditions, there is a minimal amount of drug efflux at 4 °C (Supplementary Fig 9), which increases at 37 °C. There is a significant increase in the amount of P-gp transporter (and associated activity relating to efflux of dye) under SPIR conditions at both temperatures. This effect is reversed by SPIR-W conditions at 4 °C (although not at 37 °C), after growth under SS conditions for 24 hours. In the epithelial cell line, ARPE19, there is minimal efflux of the dye in both SS and SPIR conditions at 4 °C, but increased efflux at 37 °C (only under SPIR culture conditions), with reversal by SPIR-W conditions (Supplementary Fig 9). Thus, the data clearly suggests that SPIR treatment confers a drug-efflux phenotype to cells. An improved drug-efflux phenotype is commonly thought to be a stem cell-like phenotype (Moitra *et al*., 2011).

### Factor-dependent transdifferentiation into adipocytes or osteocytes as well as differentiation into the three germ cell lineages, enabled by SPIR

We observed that SPIR transforms cells into a state in which they develop the capacity to differentiate into cells of the ectoderm, mesoderm and endoderm, by virtue of their expression of specific markers during growth in the presence of factors (Supplementary Fig 10A). Further, we observed that SPIR-treated cells are capable of transdifferentiating into adipocytes and osteocytes when the appropriate factors are added (Supplementary Fig 10B,C respectively).

### Expression of PAR2 in triple negative breast cancer (TNBC) tissue and TNBC cell line (MDA-MB231)

We examined whether PAR2 expression is elevated in cancer stemness and whether cancer cell lines respond to SPIR treatment in a manner similar to other cells and cell lines. TNBC cells are believed to be resistant to chemotherapy because of their enhanced efflux of drugs, and this is associated with cancer stemness. In the TNBC cell line, MDA-MB231, SPIR conditions resulted in increased expression of PAR2 (Fig 6A).

**Fig 6:**
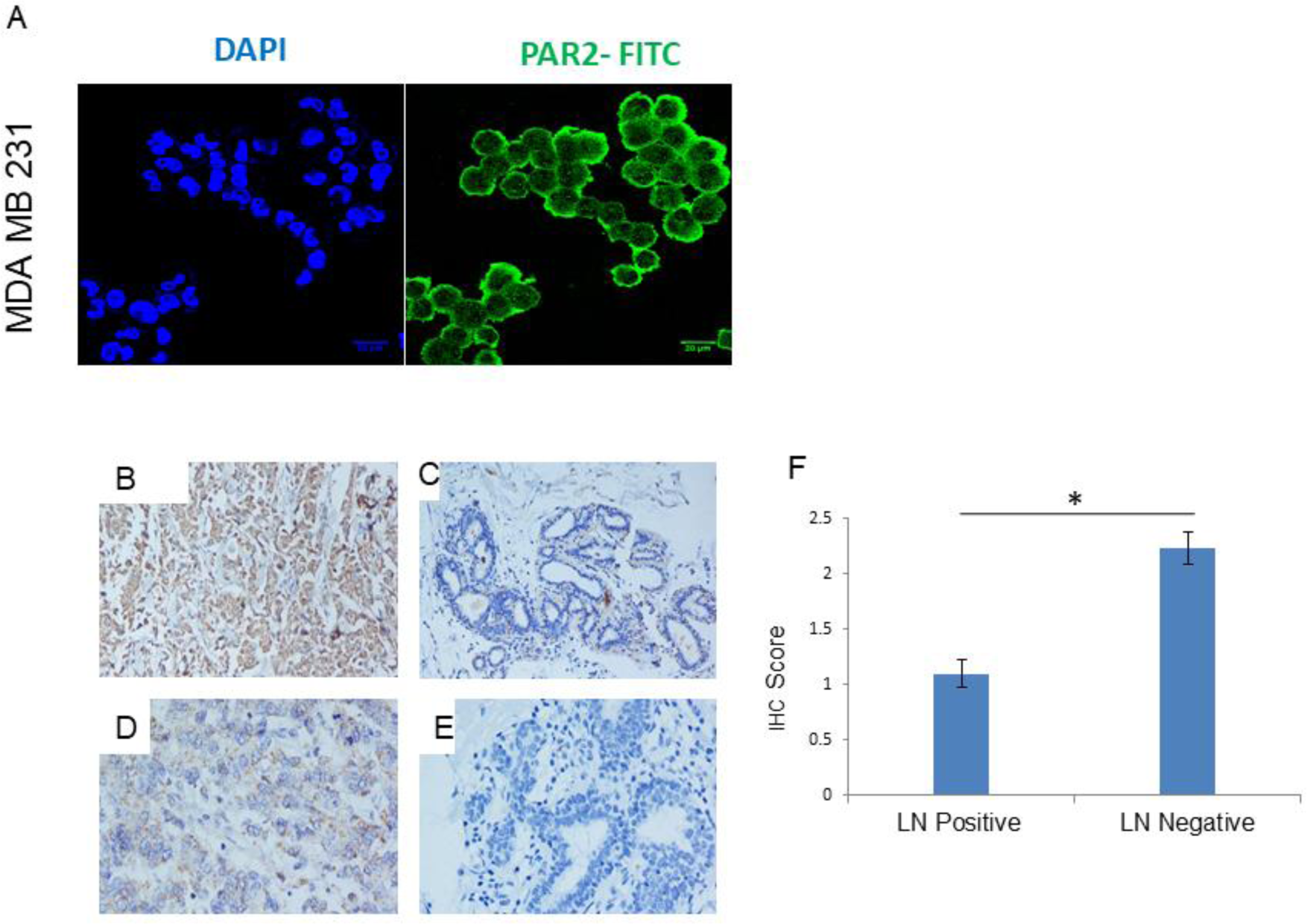
PAR2 expression observed in the TNBC cell line, MDA-MB231 under SPIR conditions (A); Representative images of PAR2 staining showing increased expression in biopsy material obtained from LN negative cases; n=30 (B), as compared to material obtained from LN positive cases; n=30 (D). In each case, adjacent, normal tissue samples were also visualized (C and E respectively). Bar diagram to compare PAR2 expression in LN negative and positive samples (F) (P value <0.0001; Mann–Whitney test).

**Fig 7:**
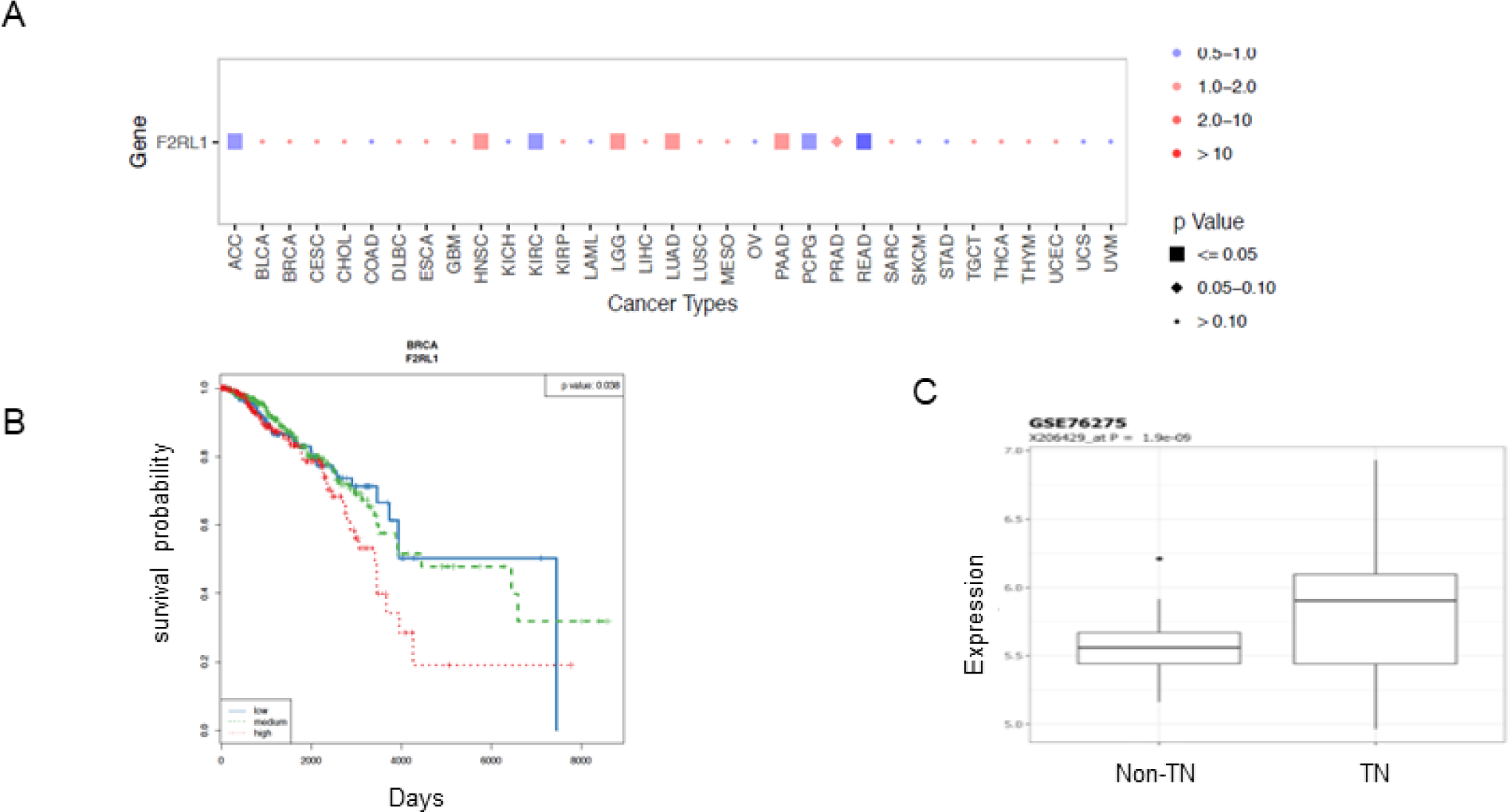
(A) Correlation of expression of F2RL1 gene (encoding PAR2) with overall patient survival plotted across all TCGA cancer types; (B) Kaplan-Meier plot showing overall survival for breast invasive carcinoma (BRCA) patients with high versus low F2RL1 expression; (C) Analysis of F2RL1 expression in publicly available human breast cancer dataset showing significantly elevated expression in triple negative breast cancer as compared with non-triple negative breast cancers.

Biopsy material was obtained from TNBC patients (patient characteristics given in Supplementary Table 2), for which staining showed consistent expression of PAR2 in all tested biopsies, irrespective of lymph node status. Notably, PAR2 expression was absent in adjacent, normal tissue sample in each case (Fig 6C, 6E). Interestingly, and apparently counter-intuitively, PAR2 expression was found to be significantly higher in biopsies derived from lymph node negative samples (Fig 6B), compared to lymph node positive samples (Fig 6D; Fig 6F) (P value <0.0001 by the Mann–Whitney test). The monoclonal antibody used for this experiment is SAM11 (Santacruz; sc13504) which we propose is specific to the epitope that binds to the tethered ligand, exposed by cleavage of PAR2 upon sustained trypsin treatment. This has been discussed in detail in a later section.

We next analyzed the correlation of expression of the F2RL1 gene (encoding PAR2) with overall patient survival, using The Cancer Genome Atlas (TCGA) resource of cancer subjects (RNASEQ data in the public domain). The survival analysis shows that higher expression of F2RL1 is associated with poor survival (P value < 0.05 and Hazard Ratio > 1) in multiple stemness-rich cancers, including head and neck cancer (HNCC), low grade glioma (LGG), lung adenocarcinoma (LUAD), pancreatic adenocarcinoma (PAAD) and prostate adenocarcinoma (PRAD) (Fig 7A). On the other hand, the under-expression of F2RL1 was associated with poor outcomes in adrenocortical carcinoma (ACC), kidney renal clear cell carcinoma (KIRC), pheochromocytoma and paraganglioma (PCPG) and rectum adenocarcinoma (READ) (Fig 7A). The Kaplan-Meier analysis on breast invasive carcinoma (BRCA) TCGA dataset suggested that higher F2RL1 expression associated with poor survival (P value <.05) (Fig 7B). Further Analysis of F2RL1 expression in a publicly available human breast cancer datasets (i.e. GSE76275) showed that the expression of this gene is significantly elevated in triple negative breast cancer (P value <1.9e-09) as compared with non-triple negative breast cancers (Fig 7C). These results indicate that F2RL1 could be a predictor of poor overall survival of breast invasive carcinoma, which could be correlated with stemness characterized by high levels of PAR2 expression.

### A discussion relating SPIR with known PAR2 signaling studies

Some insights may be gained from an analysis of what is known about PAR2 signalling. However, before examining this, it is necessary to appreciate the differences between the conditions used for the SPIR studies reported in this paper, and the PAR2 signalling studies reported by others. SPIR involves prolonged culturing of cells in trypsin (0.025 % w/v; 0.25 mg/ml; 10 μM). On the other hand, trypsin is used at much lower concentrations (5-100 nM) and over much shorter time periods (10-180 min), to examine stimulation of cells in studies involving PAR2 activation and signalling. Pathogen insult or tissue injury results in inflammatory and coagulation pathways marked by release of serine proteases like trypsin, thrombin, elastases, FVIIa, and FXa (Coughlin, 2005; Kaneider *et al*., 2007). PAR2 can be activated by several of these proteases produced during inflammatory injury including trypsin (Kaneider *et al*., 2007; Mihara *et al*., 2016). Interestingly, PAR2 is a self-activating receptor that is activated by self or neo-peptide generated following cleavage of the N-terminus of PAR2 by proteases, such as trypsin, tryptase, matriptase, factor Xa/VIIa, granzyme A, kallikreins (KLK 2/4/5/6/14), MMP-1, cathepsin S, elastase, acrosin, HAT, TMPRSS2, chitinase and bacterial gingipains (Adams *et al* 2011; Yau *et al*, 2013). Importantly, our RNASEQ data-based pathway analyses reveal that SPIR stimulates upregulation of inflammatory, coagulation and complement pathways in HCFs cultured in SPIR (e.g. C3AR1, FGB, FGG, PLAT, PLAUR, TFP1) conditions, suggesting that upregulation of these genes may be triggered following activation of PAR2. Perhaps these are the primary events triggered under trypsin culturing events, simulating injury which requires regeneration.

The canonical PAR2 cleavage site is R36S37 for serine proteases trypsin, tryptase, factor VIIa, matriptase, and few others (Yau *et al*, 2013). However, the PAR2 activation can have diverse outcomes if cleaved by other proteases like elastase and cathepsin S that target a different cleavage site, resulting in inactivation or disarming of PAR2 and generation of different tethered ligands (Adams *et al*., 2011; Ramachandran *et al*, 2012 & 2011; Elmariah 2014; Zhao *et al*., 2014a), which dictate different signaling cascades resulting in the phenomenon of biased agonism (Hollenberg *et al* 2014; Zhao *et al* 2014b). Activation of PAR2 by agonists or proteases can trigger multiple signaling pathways, such as intracellular calcium (iCa2+), MAPK, Rho kinase, nuclear factor κB (NF-κB), ERK1/2, pathways and cAMP (Suen *et al*., 2014; Dutra Oliveira *et al* 2012; Guo *et al* 2011). The activation of PAR2 results in coupling to G proteins like Gαq, Gαi, Gαs, Gα12/13 or β-arrestins to produce different downstream signaling outcomes (Ayoub *et al*., 2013, Nieman, 2016).

What could be the possible modulators functional in SPIR culturing? Our RNASEQ data from HCF cells grown under SPIR conditions shows activation of MAPK pathways (Fig 3A) and calcium signaling (HTR2A, HTR4, GNAL, ADCY7, AGTR1,CAMK2A, CYSLTR2, EDNRA, EDNRB, ITPR1, PPIF, PDE1B, PHKA1, SPHK1). PAR2 activation in monocytes leads to secretion of inflammatory cytokines IL6 and IL1β (Colognato *et al*, 2003; Johansson *et al*, 2005); interestingly, HCF cells under SPIR conditions show upregulation of many interleukins including IL6/12-like including IL6, IL11 and LIF (a stemness inducing cytokine added to stem cell media), and IL1-like cytokines including IL1α and IL1β. PARs are known to participate in diverse processes, including thrombosis, pain, and inflammation (Mrozkova *et al*, 2016; Crilly *et al*., 2012; McCulloch *et al*., 2018; Heuberger *et al*. 2019). Notably, PAR2 is being investigated as a therapeutic target for pain and inflammation and many modulators of PAR2 have been developed so far (Ramachandran *et al* 2012; Ferrell *et al*, 2008; Yau *et al*., 2013; Yau *et al* 2016; Jiang *et al*., 2018; Lee *et al*., 2019; Majewski *et al*., 2018).

Given the established roles of PAR2 in cancer, inflammation and thrombosis, how do we explain its role in cellular reprogramming in normal cells? Interestingly, emerging studies with PAR2 have begun to address the anti-inflammatory and regenerative roles of PAR2 (Piran *et al*., 2016; Rayees *et al*, 2019; He *et al*, 2018) where PAR2 promoted cell survival under stressful environment. These studies indicate role of PAR2 in tissue regeneration and homeostasis through diverse pathways; however, detailed mechanistic insights that could result in generation of stem cells upon PAR2 activation remain far from investigation. The following are some possibilities:

1. Endosomal involvement: The importance of GPCR signaling from endosomal/intracellular compartments (Hanyaloglu 2018; Weinberg et a, 2019), needs to be understood in depth so that appropriate therapeutic targets can be designed. Jimenez Vargas *et al*, (2018) demonstrated that PAR2 signaling from endosomes resulted in the persistent hyperexcitability of nociceptors (pain receptors) that mediated chronic pain in irritable bowel syndrome (IBS). Interestingly, PAR2 is hugely implicated in IBS and arthritis and both these disease pathways - IBS (IL1A, IL1B, IL18R1, IL6, STAT4, TGFB1, TGFB3) and Rheumatoid arthritis (ATP6VOB, ATP6VOD2, CTSL, IL1A, IL1B, IL11, IL6, MMP1, MMP3, TGFB1, TGFB3) are upregulated in KEGG pathway analysis of HCF SPIR cells. It indicates that PAR2 might be targeted to endosomes (e.g., via beta arrestins) with sustained signalling. Importantly, cAMP signaling events (HTR4, ATP1B1, ADCY10, ADCY7, CAMK2A, EDNRA, GIPR, PDE3A, PDE3B, PDE4D, PIK3R3, PLD1, PTGER2, SSTR2) are upregulated in SPIR; however, cAMP signaling events at the plasma membrane are usually desensitized by β-arrestin mediated delivery of phosphodiesterases, which degrade cAMP (Baillie and Houslay, 2005; Baillie, 2009). The upregulation of several phosphodiesterases (PDE3B, PDE10A, PDE4D, PDE3A, PDE1B) in HCF SPIR and upregulation of vesicular and protein trafficking pathways in RNASEQ support the contention of PAR2 endosomal signaling under SPIR conditions. The counterbalance of cAMP and beta arrestin pathways might determine the final outcome (short lived surface PAR2 signaling over sustained endosomal PAR2 signaling respectively) under SPIR conditions, which merits further investigation. Notably, secretion of bFGF and IL1β in the secretome of ARPE19 and HCF cells indicates an active secretory pathway involvement as both these cargo are signal-sequence lacking (leaderless) proteins and are known to be released by unconventional secretory pathways (Steringer *et al*., 2015; Gee *et al*, 2018); alternative secretory pathways continue to be proposed for these proteins (Monteleone *et al*., 2018; Baroza-Mazo *et al*., 2019). Interestingly, neutrophil elastase (an alternate activator of PAR2) also promotes secretion of IL1 β (Alfaidi *et al*., 2015). Whether trypsin-PAR2 mediated axis also participates in an alternative mode of secretion of IL1β and bFGF is an interesting and open question, which warrants further investigation.
2. Vesicular shedding: More importantly, trypsin-PAR2 mediated microvesicular shedding has been recently reported to aggravate triple negative breast cancer in a cell culture model of highly metastatic human breast cancer cell line, MDAMB231 through the reorganization of actomyosin network (Das *et al*, 2018 a & b). We believe that such microvesicles might underlie secretion of growth factors and cytokines in SPIR cells to promote intercellular communication and survival in SPIR cells. Notably, the phalloidin immunostainings of SPIR cells verses SS cells clearly exhibit cortical actin. It has been recently shown in the context of PAR2 activated breast cancer, that higher G/F (globular/filamentous or monomeric/polymerized) ratio for actin is indicative of a vesicle budding phenotype (Das *et al*, 2018 a).
3. Epigenetic remodeling: Epigenetic remodeling indicated by 5hmC positivity in SPIR cells might explain role of PAR2 signaling in transcriptional regulation of stem cell genes. Recently, the Tet enzyme-mediated oxidation of 5mC to 5hmC has been suggested to have a central role in epigenetic reprogramming (Hill *et al*, 2014; Finley *et al*, 2018).

At the end of this section, it may be pertinent to mention a recent study which demonstrates that collagenases as proteinases can disarm PAR2, suggesting the existence of a mechanism that suppresses PAR2-mediated inflammatory responses (Falconer *et al*., 2019). SPIR conditions with upregluated MMPs could self-limit PAR2 activity and promote its anti-inflammatory/regenerative outcomes in normal/non-cancerous cells. However, destructive role of PAR2 in tumors might be linked to sustained presence of cancer stem cell phenotype, owing to presence of redundant serine proteases that overcome MMP/collagenase mediated inactivation of PAR2.

### Background and origin of the discoveries reported here

Having described the most significant aspects of trypsin’s effects upon cells under SPIR conditions, we now describe how we chanced upon discovering these effects. While the data is detailed in the sub-sections below, a description of the chronology of our findings, and thoughts, is provided in this sub-section. The focus of the cell biological research in our group hitherto has largely been the extracellular matrix (ECM) and its various components. We have examined the role(s) of the ECM and its components in determining cell behavior in a variety of immunopathological and ophthalmological diseases. In addition to studying such behavior under ordinary culture conditions, we also use protein-engineered versions of ECM components, and examine the effects of these in comparison with non-protein-engineered components. One aspect that we commonly examine is the presence and activation status of metalloproteases such as MMP-2 and MMP-9, as a function of cell type and the presence of different factors including protein-engineered factors (Sharma *et al*., 2013; Tiwari *et al*., 2015; Tiwari *et al*., 2016; Mehta *et al*., 2018).

It is common knowledge that cells are best kept in serum-starved conditions prior to the conduct of cell-based assays, to exclude involvement of serum components. Thus, cells are maintained overnight under serum-starved (SS) conditions before any assays are carried out. It is also common knowledge that cells are unhealthy, and that some cells can die, if serum starvation is carried out for too long. Therefore, there is merit in exploring alternative culture conditions, particularly when extended culturing of cells under SS conditions is necessary, e.g., if one wants to examine an assay result as a function of time, rather than at a single time point.

In addition, since serum also affects the activation of MMPs, it is desirable to avoid serum as far as one can when one is examining MMP levels and activation. One variation to try, therefore, in respect of culture conditions is to avoid the step of first adding serum (to inactivate the trypsin) and then removing the serum (to create SS conditions, necessary for an assay), simply by avoiding the serum addition step and directly plating out cells in DMEM, without inactivating the trypsin, to see whether this affects the results (in other words, what we call SF conditions). What we found is that in the conditioned medium (CM) from such cell cultures there is a low molecular weight (LMW) band with proteolytic, and gelatinolytic, activity evident in zymograms, in a manner concomitant with greater activation of MMP-2 and MMP-9. The apparent molecular weight of this band initially led us to believe that it could be MMP-7, before we realized that it is trypsin, once we found that it is there in the zero-hour (control) time point and that it steadily reduces in intensity over multiple washings of cells through changes of the medium, suggesting that our protocol facilitated the presence of residual active trypsin (i.e., non-inactivated trypsin) from the trypsin-EDTA treatment given during passaging. This caused us to try and add trypsin exogenously to cells plated out in serum-free DMEM, and we discovered that as with the residual trypsin, this also results in progressive activation of MMP-2 and MMP-9 with time, suggesting that trypsin acts upon the zymogen forms of the said MMPS, and activates them. We then wished to see what else trypsin can do under the given conditions, i.e., when it is added to serum-free DMEM medium, or to serum-starved cells originally exposed to serum (to inactivate trypsin, after being dislodged). This led to the standardization of the SPIR conditions, and the discovery and validation of stemness. Also, we were able to conclude that non-inactivation of trypsin by serum leads to the continued presence of trypsin in cell culture, ostensibly owing to a combination of residual trypsin from the trypsin-EDTA treatment and maybe some autocrine production and secretion of trypsin by cells, which we found can be downregulated by the addition of trypsin inhibitor (aprotinin) in experiments not reported here. Together, all of these findings led us to standardize SS, SF, SFA, SPIR, SPIR-W and SPIR-F conditions.

### Presence of a low molecular weight (LMW) protease in cells cultured under SF conditions

NCI-H292 cells were cultured under SS and SF conditions for 48 hours, and zymography performed with the conditioned medium (CM) from these cultures revealed gelatinolytic activities corresponding to proteases, MMP-2 and MMP-9 (Fig 8A). CM from cells cultured under SF conditions showed greater levels of activation of pro-MMP-2 and pro-MMP-9 than cells grown under SS conditions (Fig 8B). Further, CM from cells grown under SF conditions displayed an additional band with gelatinolytic activity which was not seen in the CM from cells grown under SS conditions. The protein corresponding to this additional gelatinolytic band, unique to SF, was tentatively called LMW (low molecular weight; since it was evidently much lower in molecular weight than pro-MMP-2, pro-MMP-9, MMP-2 and MMP-9). CM from cells grown under SF conditions for ∼24 hours displayed mainly LMW, pro-MMP-2 and pro-MMP-9 (Fig 8B; lane 1); at later points of time, the intensities of the LMW, pro-MMP-2 and pro-MMP-9 gelatinolytic activities (bands) decreased, with concomitant increase in MMP-2 and MMP-9 (Fig 8B; lanes 2,3).

**Fig 8:**
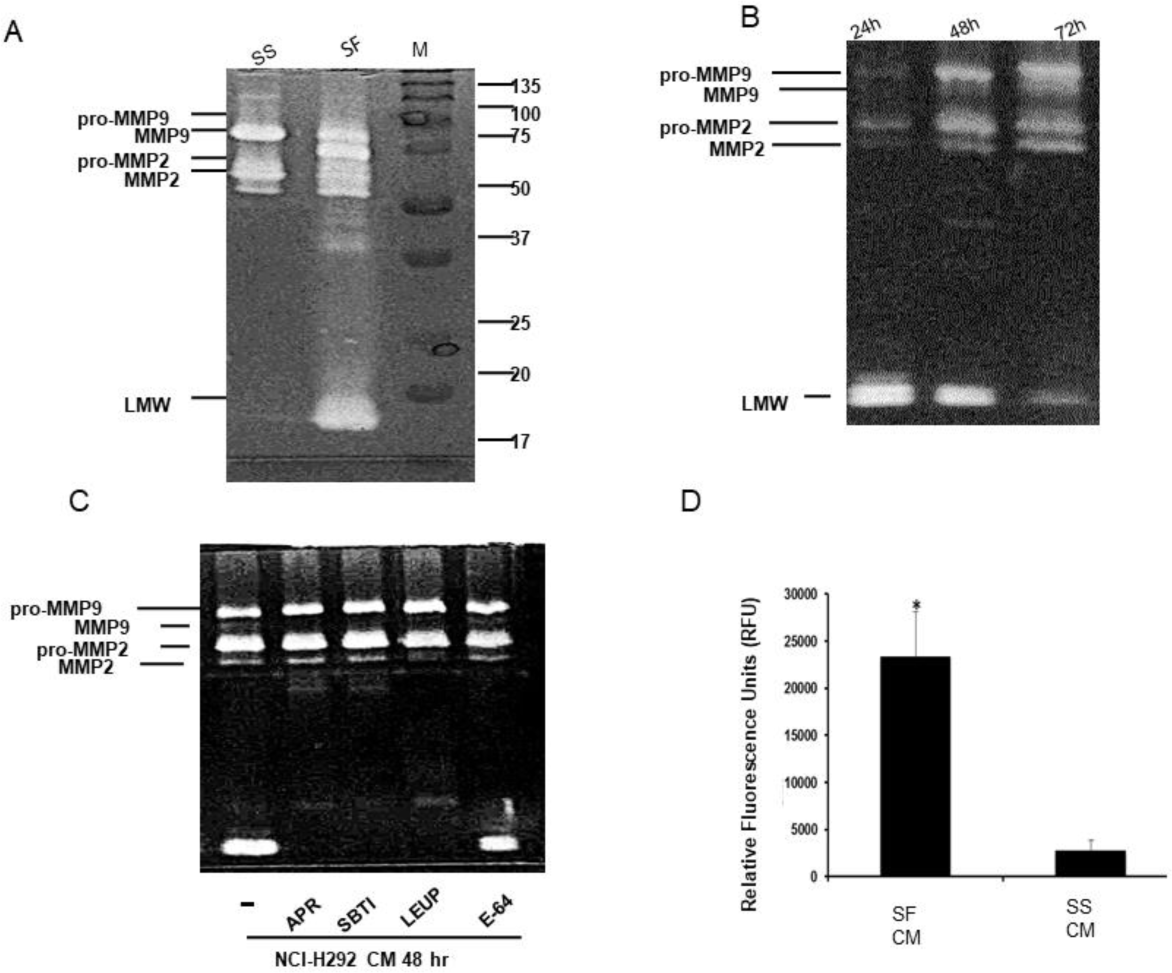
Gelatin zymogram of conditioned media (CM) of NCI-H292 cells cultured under and SS and SF conditions displayed increased activation of pro-MMP −2, −9 to their active (MMP-2 and −9) forms and a unique/novel LMW band (A); gelatinolytic profile of CM of NCI-H292 cells cultured under SF conditions over time (B Gelatin zymogram of NCI-H292 cancer cells cultured under SF conditions in presence of different protease inhibitors (C); CM of cells cultured under SF conditions can degrade MMP-specific fluorogenic substrate as detected by fluorometric assay (D).

**Fig 9:**
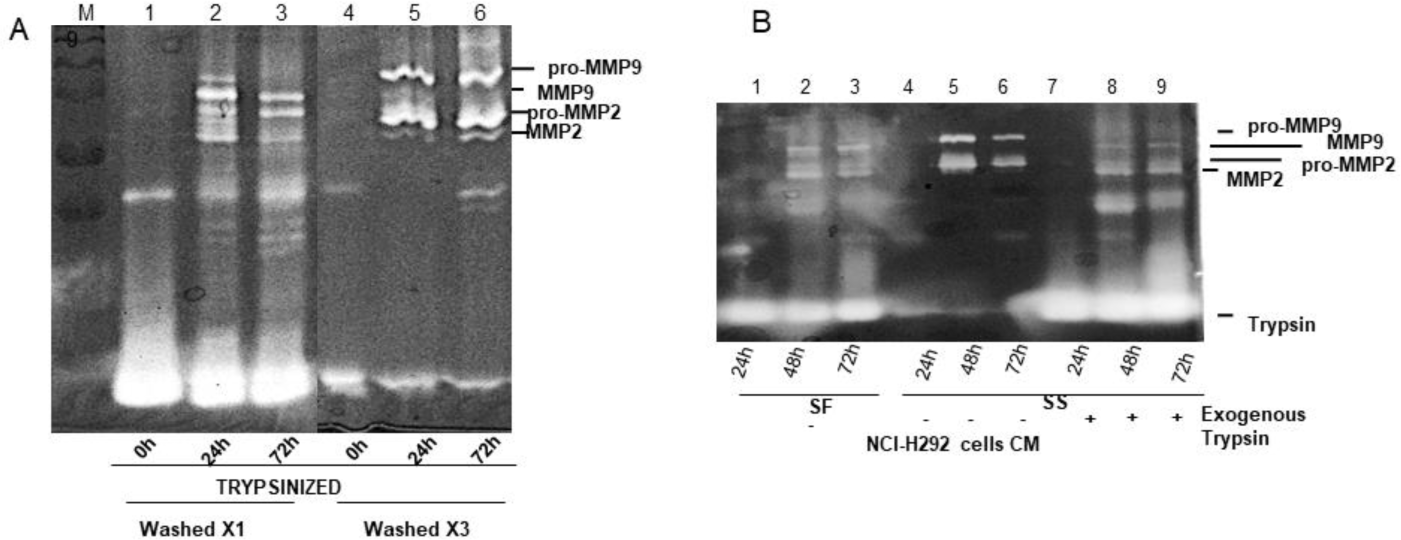
Gelatin zymogram of CM of NCI-H292 cells under SF conditions upon trypsinization, followed by either one wash or three washes with PBS (A); Gelatinolytic profile of the CM from NCI-H292 cells cultured under SF conditions (lanes 1-3); SS conditions (4-6); SS conditions + trypsin (lanes 7-9) (B).

### Apparent causal connection between the LMW activity and a serine protease

We were intrigued by the LMW band (and activity) seen in the conditioned medium from cells grown under SF, but not SS, conditions. To determine the nature of the protease associated with the LMW activity/band in the zymogram, different protease inhibitors were added to NCI-H292 cells being cultured under SF conditions, including aprotinin (serine protease inhibitor), SBTI (trypsin-like serine protease inhibitor), leupeptin (inhibitor of serine and cathepsin proteases), and E-64 (cathepsin protease inhibitor). Serine protease inhibitors were found to inhibit the LMW activity/band (Fig. 8C), indicating that LMW is a serine protease. To further explore the nature of LMW, conditioned media from cells cultured under SS and SF conditions were examined for activity against a fluorogenic peptide substrate specific to matrix metalloproteases, or MMPs (Fig 8D). Only CM from cells cultured under SF conditions favored hydrolysis of this substrate (Fig 8D), suggesting that the LMW band could potentially activate MMPs under SF conditions; however, with the same not being observed under SS conditions.

### Apparent causal connection between the LMW activity and trypsin

In the course of our experiments, we observed that (i) the LMW band exists even at the time of plating of cells (i.e., at 0 h), and (ii) the intensity of the LMW band shows variations, depending upon the number of washings given to the cells, prior to plating. We therefore cultured NCI-H292 cells under SF conditions, and examined the effects of trypsinization followed by one wash with PBS, or three washes, using zymography. Increased numbers of washes of cells after trypsinization resulted in decreased proteolytic activity corresponding to the LMW band (Fig 9A; lane 4 vs lane 1) and also in decreased intensities of the active forms of the two MMPs, MMP-2 and MMP-9 (Fig 9A; lanes 5 and 6 vs lanes 2 and 3). This indicated that residual trypsin was being washed away.

Since there was no protease other than trypsin used during the culturing of cells, the presence of the LMW proteolytic activity, even at the 0 h time point, led us to hypothesize that the LMW activity derives from the trypsin used for the passaging of cells, which is not removed prior to culture under SF conditions.

It may be noted that (a) the molecular weight of trypsin (∼24 kDa) corresponds perfectly to the gel position associated with the LMW activity (Supplementary Fig 11A); (b) trypsin is a serine protease, explaining the inactivation of the LMW activity/band by serine protease inhibitors (Fig 8C); (c) inactivation of trypsin (through addition of serum which contains anti-trypsin) is not a part of the SF conditions used by us for culturing of cells, explaining why the LMW activity is higher in the CM from cells grown under SF conditions; and (d) if LMW were indeed trypsin, this would explain why the LMW-associated activity was observed to reduce progressively with each washing of cells with PBS, as the residual trypsin left over from the initial trypsinization of cells during passaging, would be expected to reduce with repeated washings of cells with PBS, prior to plating, even if there were some endogenous production of trypsin by cells through activation of the PAR2 receptor.

This analysis further supported our hypothesis that LMW is trypsin, which correlates with our unreported finding that there are higher levels of trypsin expressed by cells in cultures under SF conditions, in comparison to SFA conditions using aprotinin (data not shown), suggesting that aprotinin does not only reduce active trypsin levels but also affects trypsin expression by cells.

### Activation of MMPs through the (necessary and sufficient) sustained presence of non-inactivated trypsin

NCI-H292 cells were grown under SF conditions, SS conditions, and SPIR conditions (Fig 1). The conditioned medium (CM) from the above cultures was evaluated for gelatinolytic activity at 24 h, 48 h and 72 h of growth. The proteolytic activity displayed by the CM from cells cultured under SF conditions proved to be entirely similar to that of cells cultured under SPIR conditions, i.e., with trypsin exogenously added to cells (Fig 9B), with a time-dependent increase in the gelatinolytic activity associated with MMP-2 and MMP-9. However, nothing even remotely similar was observed with the CM of cells cultured under SS conditions. This suggests that residual trypsin remaining over from passaging was indeed aiding in the activation of the two MMPs, and also that such activation could be induced by exogenous addition of trypsin to cells.

### Observation of the LMW gelatinolytic activity in cells other than NCI-H292

To examine whether other cells also show the MMP-activation effects of the LMW (trypsin) activity/band, we replicated the NCI-H292 experiments using other cell types, by culturing different non-cancerous cells lines under SF conditions and also examining primary cultures. Primary cultures of HCFs, and cells of the ARPE19 cell line, also showed the same phenomenon (Supplementary Fig 11B).

Thus, trypsin was established to be the key player in activation of MMPs. At the same time, as described in the beginning of the paper, trypsin was also seen to be expressed at higher levels in microarray data pertaining to cells cultured under SF conditions, in which indications of stemness were seen. Simultaneously, we found that cells cultured under SF conditions showed increased expression of stemness-associated factors, in a cytokine bead array assay. Together, all of these results pointed to a causal role for trypsin in inducing stemness, and this caused us to develop the SPIR culture conditions. Data demonstrating that SPIR conditions induce stemness has already been presented in detail in the earliest sections of this paper.

## DISCUSSION AND CONCLUSIONS

We believe that trypsin-mediated activation of the PAR2 receptor could trigger multiple events, which remain to be fully delineated. We are working to elucidate these. We think, for example, that trypsin could provide a signal for further expression and secretion of trypsin (to activate an autocrine mechanism utilizing trypsin-signaling), which can be downregulated by trypsin inhibitors. There is evidence that trypsin is secreted at concentrations compatible with activation of PAR-2 by colon cancer cell lines (Ducroc *et al*., 2002), indicating that trypsin exerts an autocrine/paracrine regulation of PAR-2 activity (Darmoul *et al*., 2001). Others have also reported autocrine secretion of trypsin in pancreatic cancer (Shimamoto *et al*., 2004), esophageal adenocarcinoma (Han *et al*., 2013) and breast cancer cell line MDA MB-231 (Ge *et al*., 2004). We argue that in the presence of aprotinin, trypsin is inactivated, thus making it unable to provide a positive feedback for further expression of trypsin. This autocrine expression of trypsin needs further experimentation in the SPIR-based system. So does the expression of factors such as bFGF and IL1β. Besides this, many other aspects remain to be explored, beyond the ones described here.

The main point of importance, of course, is the observation that after SPIR treatment, rounded-off cells cluster together and become enabled to either undergoing transdifferentiation or differentiation into any of the primary cell lineages (ectoderm, endoderm, or mesoderm). The SPIR culture conditions appear to be uniquely favorable for the generation of stemness characteristics: (i) since the SPIR treatment (with trypsin) is done under nearly 80% confluent conditions, and the frequency of conversion of cells to stem cell-like cells appears to be significant, this method can overcome the limitations of low efficiency and incomplete reprogramming associated with the conventional methods of generation of iPSCs; the previously known methods of reprogramming are plagued by low efficiency ranging from around 4% using modified messenger ribonucleic acid, to 1% with integrating viruses, and to 0.001% with the non-integrating Sendai virus, adenovirus, plasmids and direct protein delivery (Zhou *et al*., 2013; Iseki *et al*., 2016; Bar-Nur *et al*., 2014; Buganim *et al*., 2014; Hou *et al*., 2013; Carey *et al*., 2011; Warren *et al*., 2010; Warren *et al*., 2012).; (ii) since SPIR is very simple and uses minimal reagents after serum starvation done in routine culture conditions, it offers robust experimental reproducibility; (iii) the simplicity of the procedure is coupled with less hands-on time and low costs for reprogramming; (iv) the absence of any retroviral vectors adds the much needed safety features (e.g., possible avoidance of teratoma formation) for potential use of this technology in regenerative medicine in the future; (v) the SPIR method of reprogramming is not limited to fibroblasts, since we have shown stem cell-like characteristics in different kinds of source/parent cells (including epithelial cell line, primary culture of fibroblasts, cancer cell lines etc); however, according to the RNA sequencing data, the SPIR reprogrammed cells showed better enrichment scores when primary cultures of fibroblasts (HCF) were used, compared to what is seen with epithelial cells (ARPE-19).

We have shown that SPIR-reprogrammed cells have the key features of their physiological embryonic stem cell counterparts. These include cell cycle arrest, expression of stemness markers (Oct4, Sox-2, Nanog) along with ki67, and increased expression of bFGF and IL-1β. IL-1β is known to promote expression of stemness markers and drug resistance, and bFGF plays a crucial role in sustenance of self-renewal and pluripotency of hESCs (Greber *et al*., 2010; Wang *et al*., 2005; Xu *et al*., 2005; Lanner and Rossant, 2010). The serine protease-mediated (trypsin-mediated) reprograming of cells was confirmed by the RNA-sequencing data, illustrating the induction of stemness through pathways known to be associated with stemness (Fig 3A), such as the Notch pathway (Liu *et al*., 2010), TGF-β (Wang *et al*., 2019), IL-6 (Brady *et al*., 2013), p38 MAPK (Mitra *et al*., 2018) and NRF-2 (Ishii *et al*. 2014, Chang *et al*., 2018; Diehl *et al*.. 2018; Hawkins *et al*., 2016). It is to be noted that the TGFβ pathway is important for pluripotency due to its direct modulation of expression of Nanog (Xu *et al*., 2008). The increased expression of genes related to the glycolytic pathway in ARPE-19 cells, upon SPIR treatment, reflects the pluripotent state of these cells under these conditions, since aerobic glycolysis supports the maintenance of the pluripotent state by maintaining histone acetylation and the associated open chromatin structure of PSCs (Moussaieff *et al*., 2015; Gu *et al*., 2016; Folmes *et al*., 2011; Guda *et al*., 2018; Cliff *et al*., 2017, Dahan *et al*., 2019).

An examination of Table 2 shows that under SPIR conditions whereas the primary culture of human corneal fibroblasts (HCF) favors expression of genes associated with generation of hematopoetic and mammary stem cells, the ARPE-19 cells favor the formation of neural stem cells and embryonic stem cells. The generation of neural stem cells in the latter can be justified because retinal pigment epithelial cells are originally derived from the neuro ectoderm (Garcia-Ramirez *et al*., 2011); the generation of embryonic stem cells can be attributed to the abundant amounts of bFGF produced by these cells under the SPIR conditions (Table 1). It is well known that FGF signaling plays an important role in the self-renewal property of human ES cells (Levenstein *et al*., 2006). It has been shown that in the absence of fibroblasts, increased amounts of FGF2 (40 ng/ml) can support the growth and maintenance of human ES cells (Wang *et al*., 2005; Liu *et al*., 2006; Xu *et al*., 2005). The characteristic feature of SPIR-treated ARPE-19 cells is the abundant formation of bFGF (∼10ng/ml), which can explain the bias towards formation of ES cells. This high abundance of bFGF levels in the CM of ARPE-19 cells is potentially useful as a stem cell maintenance medium, since sustained levels of bFGF improve the maintenance of human pluripotent and neural stem cells (Lotz *et al*., 2013). There have been persistent efforts to increase bFGF bioavailability (Han *et al*., 2017; Dvorak *et al*., 2018; Horighuchi *et al*., 2018; Titmarsh *et al*., 2017) in stem cell reserach because it has low thermal stability, being highly labile at 37°C (Levenstein *et al*., 2006; Furue *et al*., 2008), causing bFGF production efforts to suffer from batch-to-batch variations, and high costs of production (Ueki *et al*., 2019). In light of this, our finding regarding the excessive production of bFGF by a certain type of SPIR-treated cell (ARPE-19) assumes great importance, since the conditioned medium from SPIR treatment of ARPE-19 could support the maintenance of undifferentiated cells in all kinds of experiments, in the future.

Interestingly, in the course of studying co-localization of PAR2 and stem cells markers, we needed to use PAR2 antibody from a host other than mouse (rabbit monoclonal to PAR2: Abcam-180953) since the antibodies for the stem cell markers available with us were in the mouse background. To our surprise, we found that the rabbit antibody to PAR2 did not distinguish between the SPIR and SS treated cells (data not shown), unlike the mouse PAR2 antibody (mouse monoclonal to PAR2: Santacruz-SAM11-sc-13504). An examination into the reason for this observation led us to the possibility that although the mouse PAR2 antibody is directed against the epitope comprised of amino acids 37-50 of PAR2 of human origin (which happens to also be the site for the tethered ligand, resulting in activation of PAR2 receptor), the rabbit monoclonal antibody, on the other hand, is directed against an epitope towards the C-terminus of PAR2 (exact epitope is proprietary). Therefore, it is possible that the mouse monoclonal antibody can differentiate between the inactive and active states of PAR2, which is seen in SS and SPIR treated cells respectively, using the mouse monoclonal antibody only. We propose that the trypsin in SPIR cultures activates the PAR2 by generating the tethered ligand for the receptor. This could also explain the unexpected staining of PAR2 observed in cells concomitant with differentiation into cells of different lineages under SPIR conditions (Supplementary Fig 10); here too, we needed to use the rabbit monoclonal antibody for PAR2 staining since the lineage marker antibodies were in the mouse background. We remain open to the possibility that there is spatial and/or temporal expression of PAR2, which needs further experimentation. In addition, co-localization of PAR2 with stem cell markers remains to be assessed.

Further, the role played by PAR2 in the stemness generating process is suggested by the fact that all triple negative breast cancer patient biopsies exhibited expression of PAR2, with lymph node negative samples showing statistically significant increase in expression, in comparison with biopsies from lymph node positive cases. This is in line with previous studies which showed that the breast cancer stem cells (CD44+/CD24−/low tumor cells) are inversely associated with lymph node metastasis (Mylona *et al*., 2008). Also, it has been shown that CD44−/CD24+ tumor cells are associated with a worse clinical outcome (Baumann *et al*., 2005), since CD24 expression causes cancer cells to acquire characteristics which aid metastasis (such as increased migration and invasion). This would signify that expression of PAR2 could serve as an early marker of TNBC which, upon differentiation, may promote metastasis. This finding is important in the context of Triple-negative breast cancer (TNBC). This is because of the TNBC subtype being negative for estrogen receptor (ER), progesterone receptor (PR), and human epidermal growth factor receptor-2 (HER2); there is no option of endocrine or any other targeted therapy. Given that the TNBC subtype is very aggressive and prone to development of resistance to cytotoxic chemotherapy with chances of early relapse, it is imperative that novel biomarkers for targeted therapy and prognosis are developed. In light of our data, PAR2 seems like a promising target for therapy, and since registered, FDA-approved drugs have been seen to act as PAR2 antagonists (Xu *et al*., 2015), use of such re-purposed drugs for PAR2 would be a viable option for the development of TNBC treatment(s).

To evaluate whether expression of the gene corresponding to PAR2, i.e., F2RL1, correlates with poor prognosis, we performed Kaplan-Meier test analysis of overall survival. Our results showed that patients with increased expression of F2RL1 gene had a worse 5-year overall survival than patients with decreased expression (*p* = 0.03). This means that increased expression of PAR2 in lymph node negative cases (as seen by us), would imply an increase in “stemness” or presence of poorly differentiated tumor cells in lymph node negative cases, which going by the Kaplan-Meier analysis results, would suggest a trend towards worse 5-year overall survival. We propose that the expression of F2RL1 could serve as a potential prognostic marker.

## CONCLUSIONS AND PERSPECTIVES

This work began with the serendipitous discovery that trypsin in conditioned medium is associated with higher levels of MMP-activation. It went on to demonstrate that sustained exposure to trypsin which has not been inactivated by A1AT/serum causes cells in DMEM to attain and maintain a state of stemness, with apparent involvement of the PAR2 receptor, the levels of which are increased with trypsin exposure.

This is certainly not the first time that cells have been exposed to trypsin in cell culture. Nor is it the first time that trypsin has been linked to cell health. However, it is probably the first time that cells have been exposed to trypsin over such extended time periods, and then analyzed, explaining why the stemness induced by trypsin has previously escaped the notice of other researchers.

Of course, it is also possible that some of the effects of extended exposure to trypsin have been noticed before, and that this was, in fact, responsible for the very origins of the practice of using serum to inactivate trypsin before re-plating of cells. In other words, it is possible that researchers discovered very early during the development of cell culture methods that cells do not change shape or morphology during very brief exposures to trypsin-EDTA, but that over extended periods of exposure cells do show changes (which are, fortunately, or so it was probably thought, eliminated through the addition of serum). During the beginnings of the development of cell culture methods, it would have been impossible for researchers to recognize that the rounded, agglomerating and clustering cells that result from extended exposure of cells to trypsin possess some kind of stemness, and that such cells are not ‘unhealthy’ cells.

Therefore, we feel that we are truly fortunate that the underlying science which was necessary to understand the observations presented in this paper were already in existence, before our observations, due to the gradual maturation (and coalescence) of the sciences of developmental biology, cell biology, molecular biology and biochemistry. These fields had neither matured, nor come together, when the practice of adding serum to inactivate trypsin must have been initiated. At the same time, given the indoctrination of always using serum to inactivate trypsin, we were fortunate to have chosen to not bother to inactivate trypsin by serum, to see the effects. As is often said, ‘sometimes fools rush in where angels fear to tread’ and it is possible that we have been such fools, with some happy consequences. Of course, there is merit in doing experiments that lie outside the box, and the points in any data that are outliers often have the greatest potential for pointing to new phenomena.

It is interesting that although trypsin is made in the pancreas and used in the intestine, and alpha-1 anti-trypsin (A1AT) is made in the liver, both trypsin and A1AT are also present in the serum at significantly high levels (Hendstrom *et al*., 1996; Donato *et al*., 2012). Presumably trypsin and A1AT exist in some sort of delicate balance, and the existence of such a balance has been proposed (Pickrell, 1987), in respect of mechanisms explaining why deficiencies, as well as excesses, in the amounts of either of these proteins lead to different diseases (Qing-quan *et al*., 2013). There is also evidence suggesting that trypsin is associated with cell proliferation (Ohta, 2003) and with certain cancers (Darmoul *et al*., 2001).

Trypsin appears to be present in the serum to fulfil some important purpose(s) and clearly the presence of trypsin and A1AT in the serum are somehow important to the organism, since their lack or excess cause diseases. However, our perspective is that if trypsin and A1AT were only important for certain organs and/or for certain tissues, organisms could easily have evolved to produce and distribute these substances locally, since it often seen that the same protein is synthesized and secreted by different cell types separated in space as well as time. The high levels of presence of trypsin and A1AT in a fluid such as the serum, which bathes all cells in the entire organism, suggests that trypsin and A1AT have a higher function, which has not been described before, i.e., a function involving all cells and tissues. The fact that there is a trypsin-specific receptor like PAR2 which exists at some basal level of expression on all cells, with a feedback mechanism ensuring upregulation in the presence of active trypsin (as is already known) also suggests that there is a higher function of trypsin which happens to service all cells, presumably whenever it is required. We believe that we have found this function. It is that sustained exposure to active trypsin, with no inhibitors (or low levels of inhibitors, ineffective in inactivating the entire trypsin present in the environment) determines cellular regeneration potential.

The organism needs to be able to regenerate certain parts of itself, at certain points of time. Gurdon showed that the nuclei of most adult cells can be persuaded to reprogram themselves in the environment of the egg cell’s cytoplasm, to regenerate an entire organism, through the attainment of totipotence (Gurdon 1962). Yamanaka showed that it is possible to persuade cells to approach a pluripotent stem cell state by expressing four specific factors within such cells, and then allowing intracellular cell signaling and cascades of gene expression to occur, to reprogram nuclei sufficiently to generate cells of all three lineages, including ectoderm, endoderm and mesoderm, but not the trophectoderm (Takahashi and Yamanaka, 2006).

Gurdon’s reprogrammed cells give rise to an entire organism, but they have lost the ability to go back to being the original cell type they were, to begin with, through any kind of de-induction. The same can be said of the reprogrammed iPSCs obtained by the introduction of the Yamanaka factors, which can also give rise to all cells barring trophectodermal cells, but which do not naturally go back to becoming the original cell type through any kind of de-induction. In contrast, the cells we describe here have characteristics of trypsin-induced stemness, and have also clearly undergone reprogramming of some sort, in their nuclei; however, the level of the reprogramming is also clearly much softer, since it is reversible. While maintenance of trypsin in the medium (DMEM) makes cells ready to be reprogrammed into any of several different types of cells (and, of course, this needs to be further explored by many more investigators), it is much more interesting that cells go back to being what they were originally, and do not remain stem cells, when trypsin is withdrawn. This sort of stemness, in which cells have been only partially re-programmed without forgetting their origins, has not been reported before.

We would thus appear to have discovered a form of stemness which is suited to cellular transdifferentiation, which is probably what the fully-grown organism requires for minor repairs and regeneration in place of the irreversibly-induced stemness of the iPSCs which can give rise to cancers if cells do not find the correct factors and conditions for differentiation, in the appropriate environments. In trypsin-induced stemness, cells appear to differentiate back into the stemness-progenitor cell type when factors are not available, and there appears to be less scope for uncontrolled growth and division. Instead, due to the inhibition of trypsin by A1AT in the serum, it would appear that trypsin-induced stemness arises only when there is a transient and local reduction in A1AT, or increase in trypsin availability (overwhelming the inhibitory effect of A1AT) together with feedback-based upregulation and activation of PAR2, e.g., at the sites of injuries, where healing and regeneration are required.

## ACKNOWLEDGEMENTS

MLG acknowledges intramural grants (No. 71/7/Edu-14/275 and 71/2-Edu-16/4840) from the Postgraduate Institute of Medical Education and Research (PGIMER). PG acknowledges intramural core funding from the Indian Institute of Science Education and Research (IISER) Mohali, as well as funding from a general grant (MHRD-14-0064) for a Centre of Excellence in Protein Science, Design and Engineering (CPSDE) from the Ministry of Human Resource Development (MHRD), Govt. of India.

## AUTHORS’ CONTRIBUTIONS

MS and MLG discovered the effect, invented SPIR, and were involved in the planning, design, execution, supervision and analysis of all primary and secondary data for all experiments, as well as in the coordination of all studies and integration of all the data into a cogent hypothesis/concept and in the writing of the manuscript. PG joined the study during the discovery of the effects of exogenous addition of trypsin, and thereafter participated in all discussions, supervision of some experiments involving immunofluorescence, cytometry, and morphology, and was also involved in some coordination of different studies and in certain parts of the creation of the hypothesis/concept and helped in writing and editing of the manuscript. RK was involved in the confocal immunofluorescence experiments confirming earlier immunofluorescence observations, in cytometry experiments, and in the microscopic observations of morphological changes. SS, BT and MB were involved in generating and analyzing all RNASEQ data, based on cellular RNA purified, quality-checked and supplied by MS and MLG, and also in all comparative transcriptomic analyses, including comparative analyses of experimental SPIR data with stem cell data and differentiated cell data available from databases, as well as in the bioinformatics analyses of PAR2 gene expression in cancers, all of which was performed under MB’s supervision. MS and MLG also participated in detailed comparisons of RNASEQ results with the literature. GS performed surgery and provided tissue samples from TNBC patients. GK and AB analyzed the IHC studies of biopsies of tissues from TNBC patients. JR performed surgery and provided ocular tissue from which primary cultures of HCFs were established. All human tissue samples were handled in PGIMER under institutional ethical clearance up to the point of creation of microscopic slides, after fixing.

---

**Fig S1:**
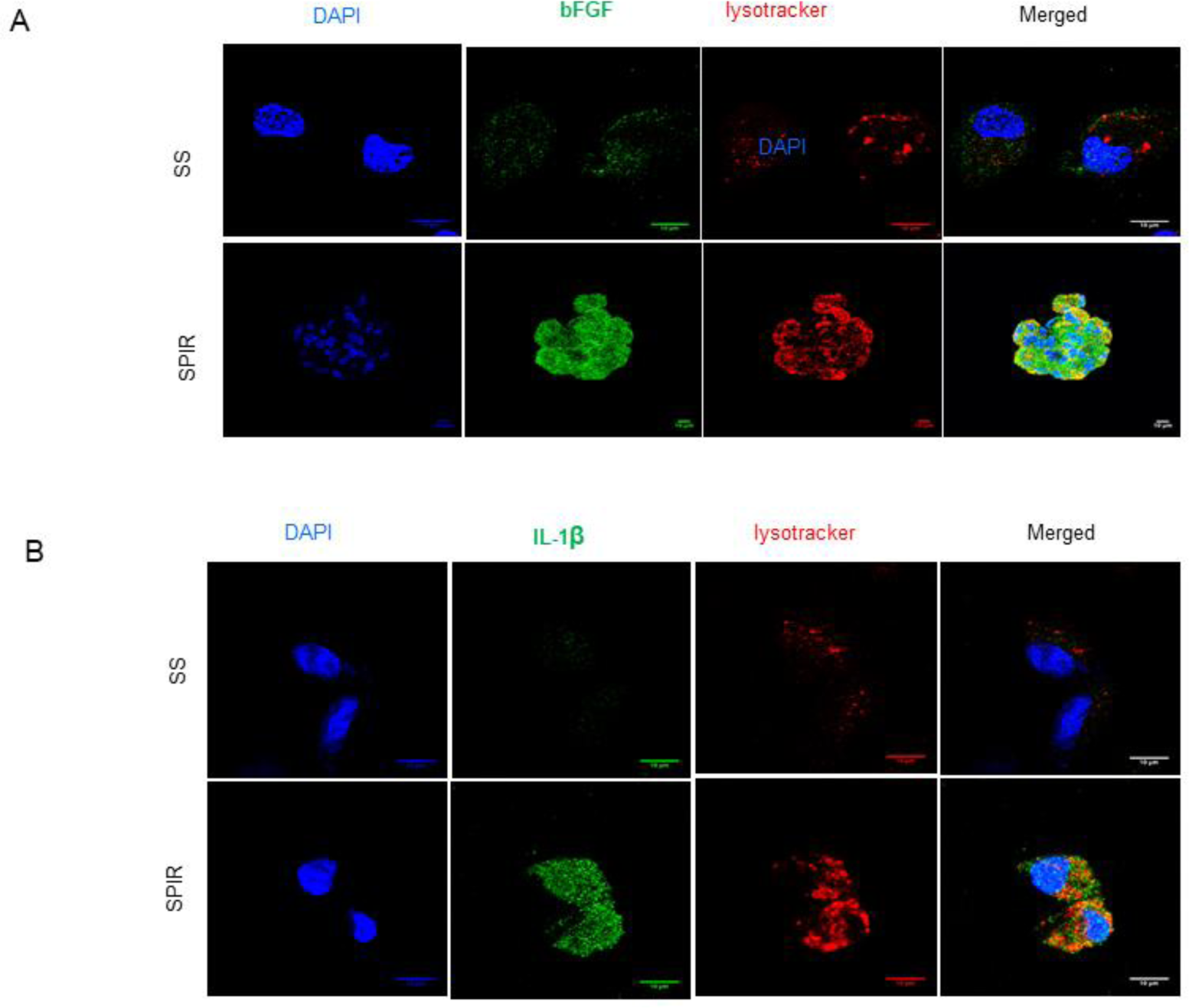
Immunostaining of ARPE-19 cells for bFGF (A) and IL-1β (B) (FITC-green), DAPI nuclear staining (blue), lysotracker dye (red), along with merged fluorescent images captured by confocal microscopy (Scale bar = 10 μm).

**Fig S2:**
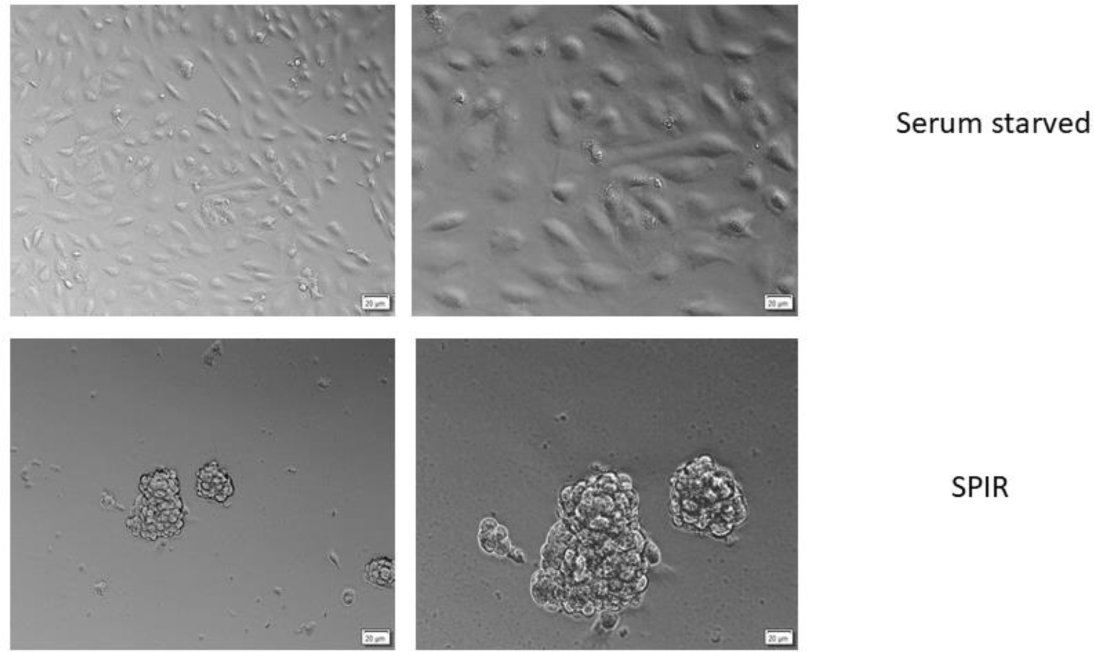
Morphology of ARPE-19 cells under serum starved (SS) and SPIR conditions.

**Fig S3:**
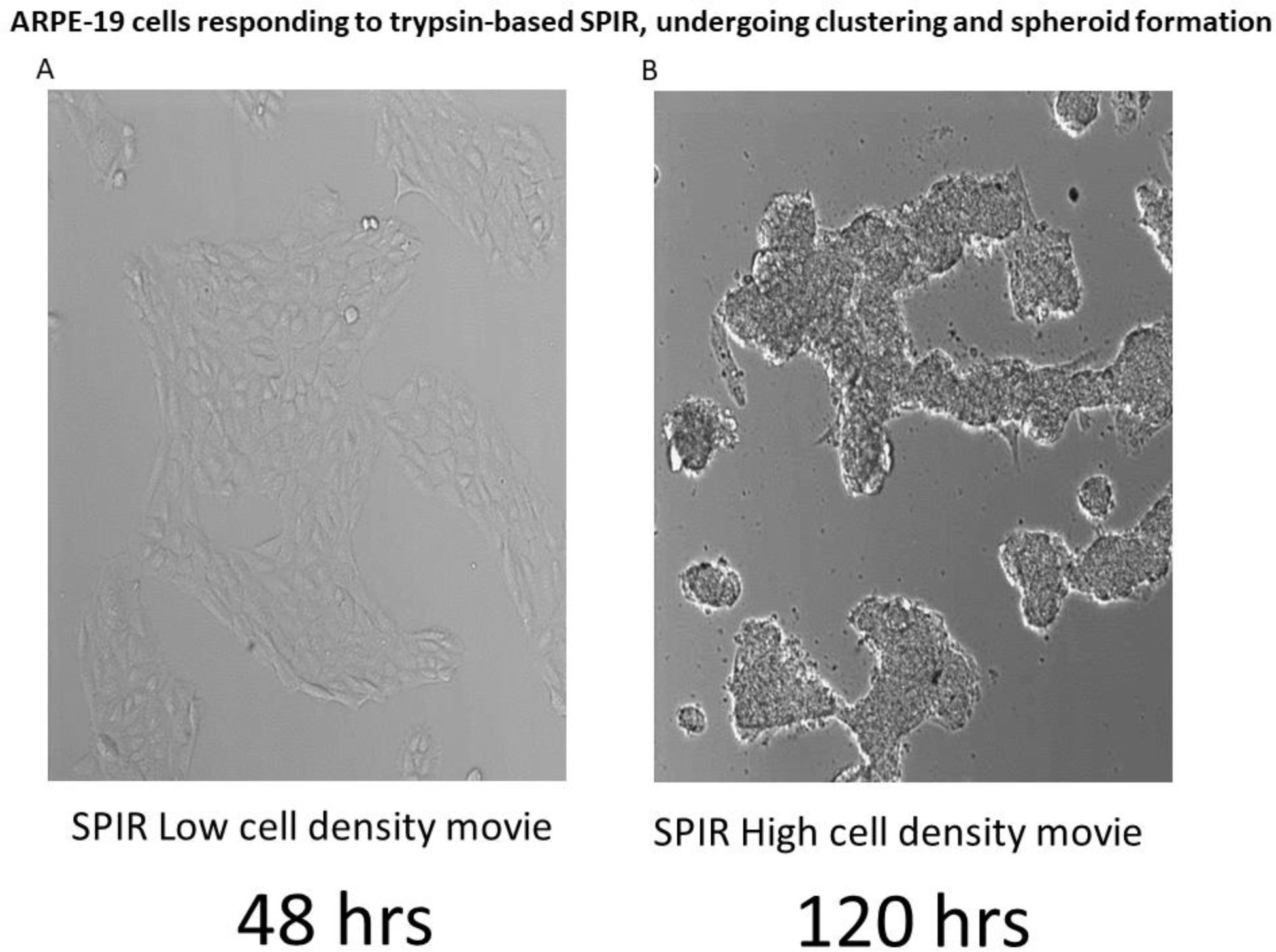
Changes in morphologies of ARPE-19 cells over time as they are subjected to SPIR. The cells become rounded, undergo clustering, and form spheroids. Representative data is provided in this supplementary figure which actually contains a pair of movies in which live cells are tracked over time on a widefield microscope. Copy and use the googledrive link (https://drive.google.com/open?id=1gPl_3IMAQ4Bz4306K2NfbVsNw6ZCF2r2) to access and download the movie file(s), which are embedded in an MS Powerpoint slide, and can run within the Powerpoint App. (A) SPIR treatment of cells growing at low density, (B) SPIR treatment of cells growing at high density.

**Fig S4:**
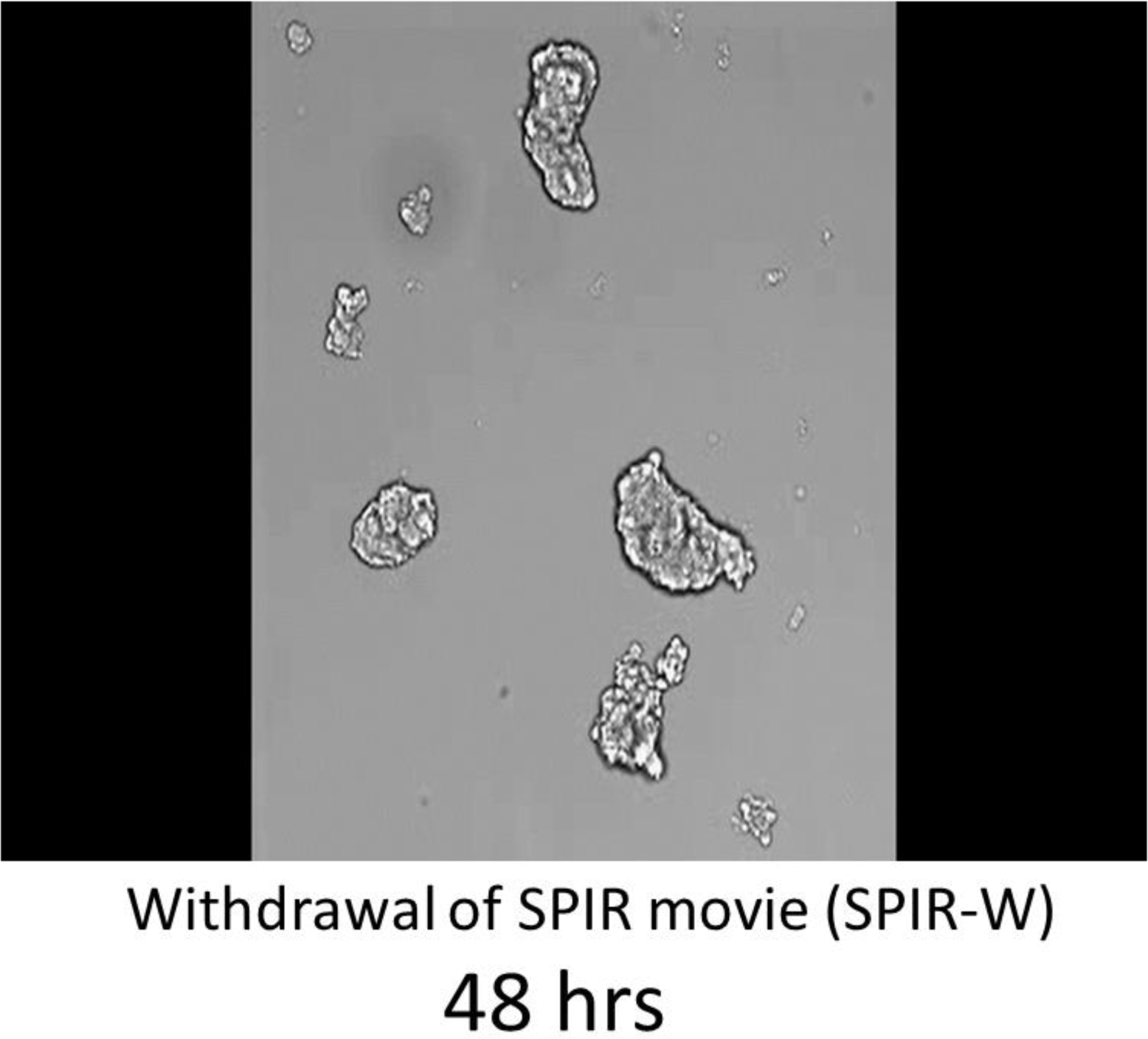
Changes in morphologies of ARPE-19 cells over time as SPIR conditions are withdrawn. The cells lose their rounded shapes, undergo de-clustering from their spheroid states, and return to the ARPE-19 morphology. Representative data is provided in this supplementary figure which actually contains a movie in which live cells are tracked over time on a widefield microscope. Copy and use the googledrive link to access and download the movie file, (https://drive.google.com/open?id=1ll5wvBPVDizFJI3JH8RhhEYjayBxFjGu) which is embedded in an MS Powerpoint slide, and can run within the Powerpoint App.

**Fig S5:**
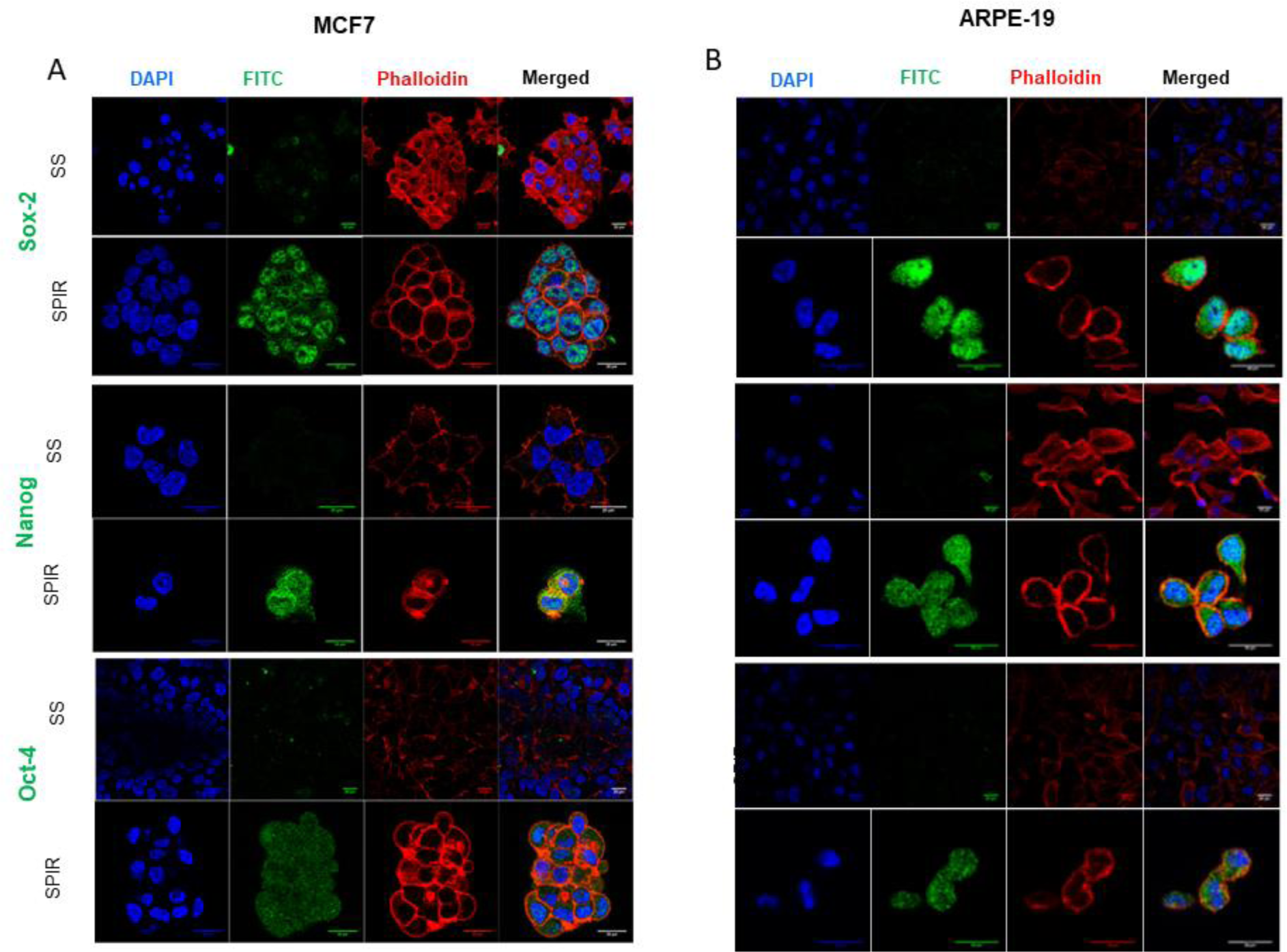
MCF7 (A) and ARPE-19 cells (B) were cultured under SS and SPIR conditions. TRITC-Phalloidin labeled F-actin (red), FITC-labeled pluripotency markers (green), DAPI nuclear staining (blue) and merged fluorescent images were captured by confocal microscopy (scale bar = 20 μm).

**Fig S6:**
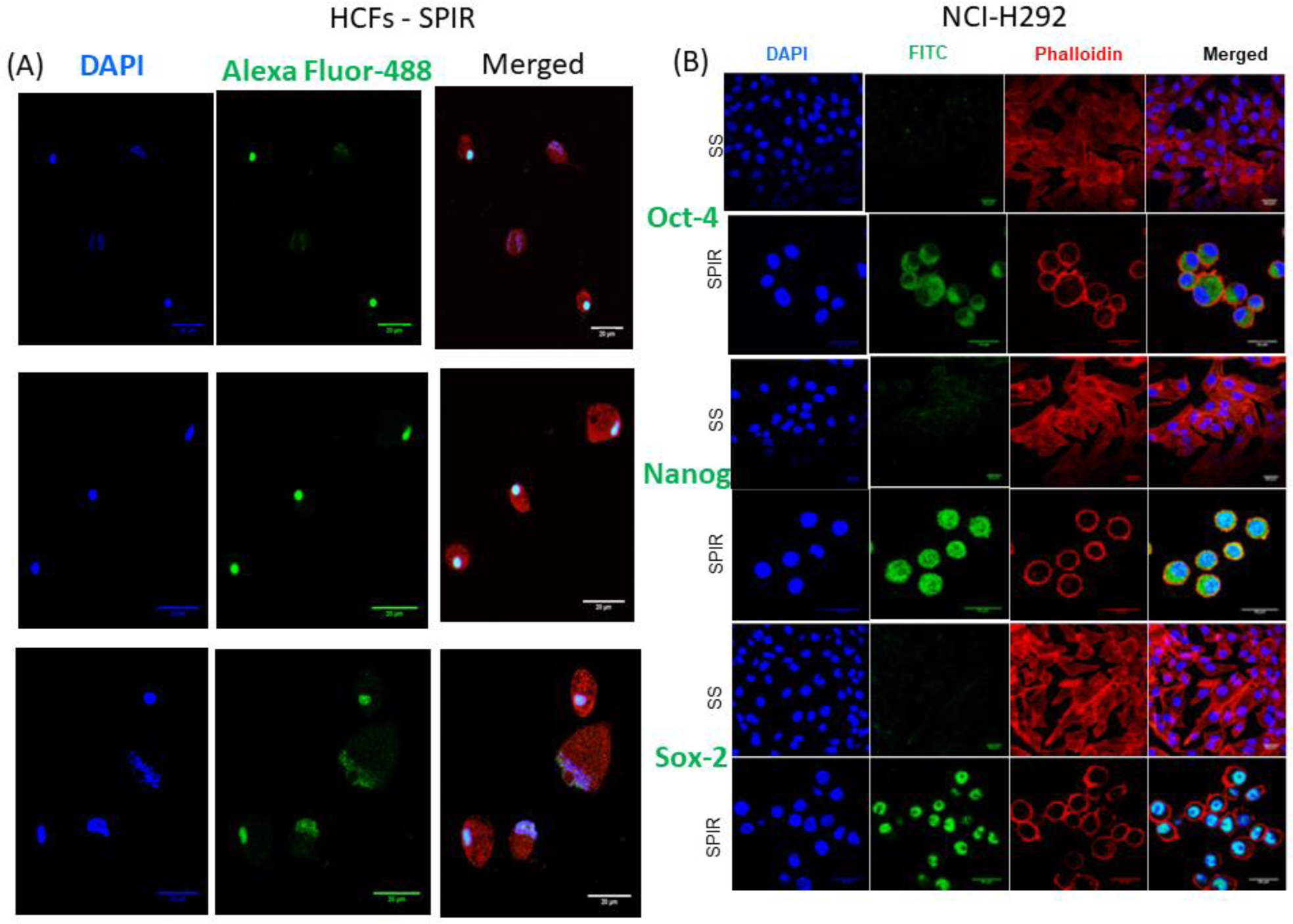
Immunostaining of primary culture of Human corneal fibroblasts (HCF) (A) and the mucoepidermoid pulmonary carcinoma cell line (NCI-H292 cells) (B) for TRITC-Phalloidin labeled F-actin (red), FITC-labeled pluripotency markers (green), DAPI nuclear staining (blue) and merged fluorescent images, captured by confocal microscopy (scale bar = 20 μm).

**Fig S7:**
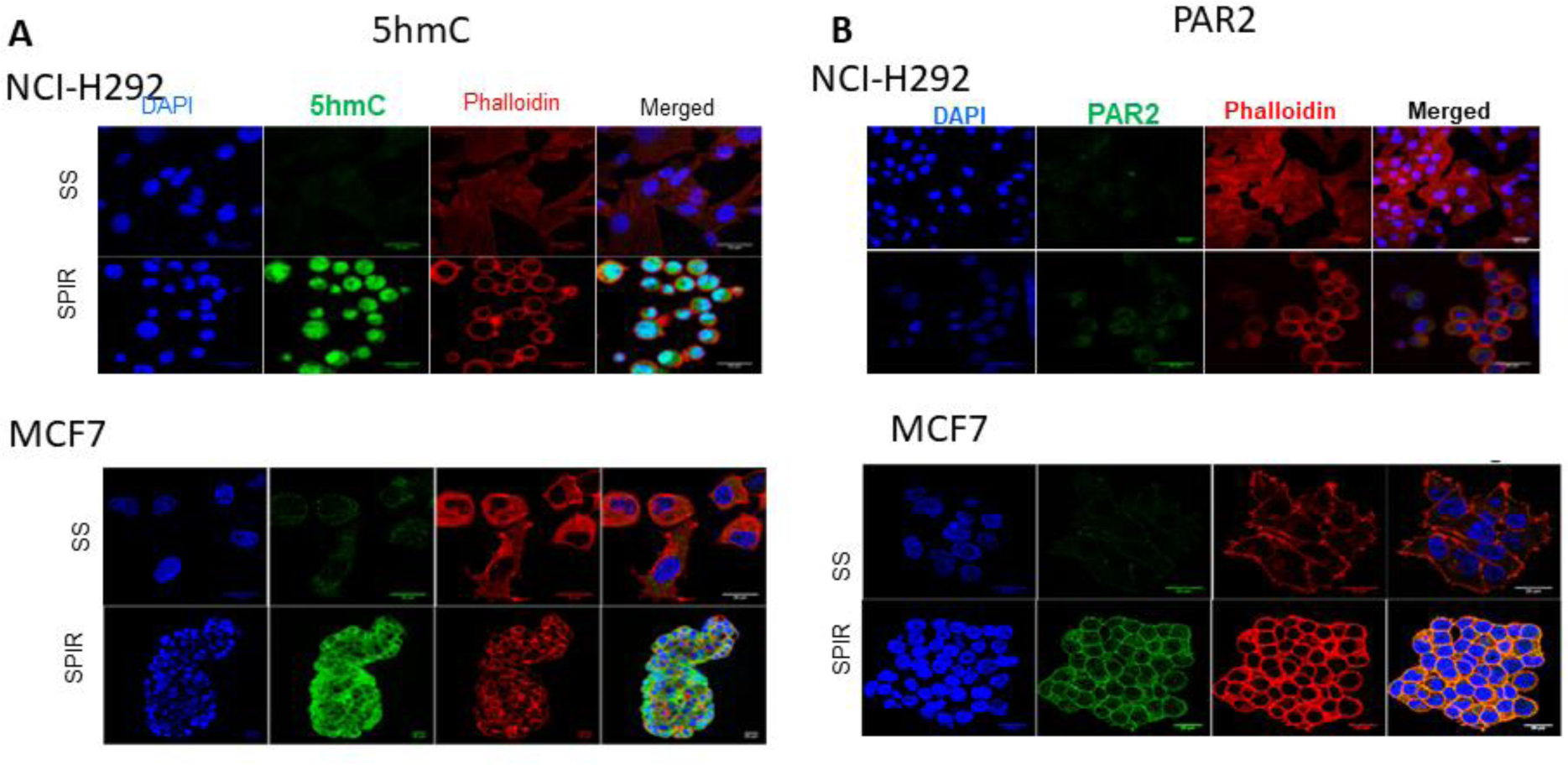
Immunostaining of NCI-H292 cells and MCF7 cells for DAPI nuclear staining (blue),TRITC-Phalloidin labeled F-actin (red), FITC-labeled (green) for 5hmC (A) and PAR2 receptor expression (B), along with merged fluorescent images, captured by confocal microscopy (scale bar = 20 μm).

**Fig S8:**
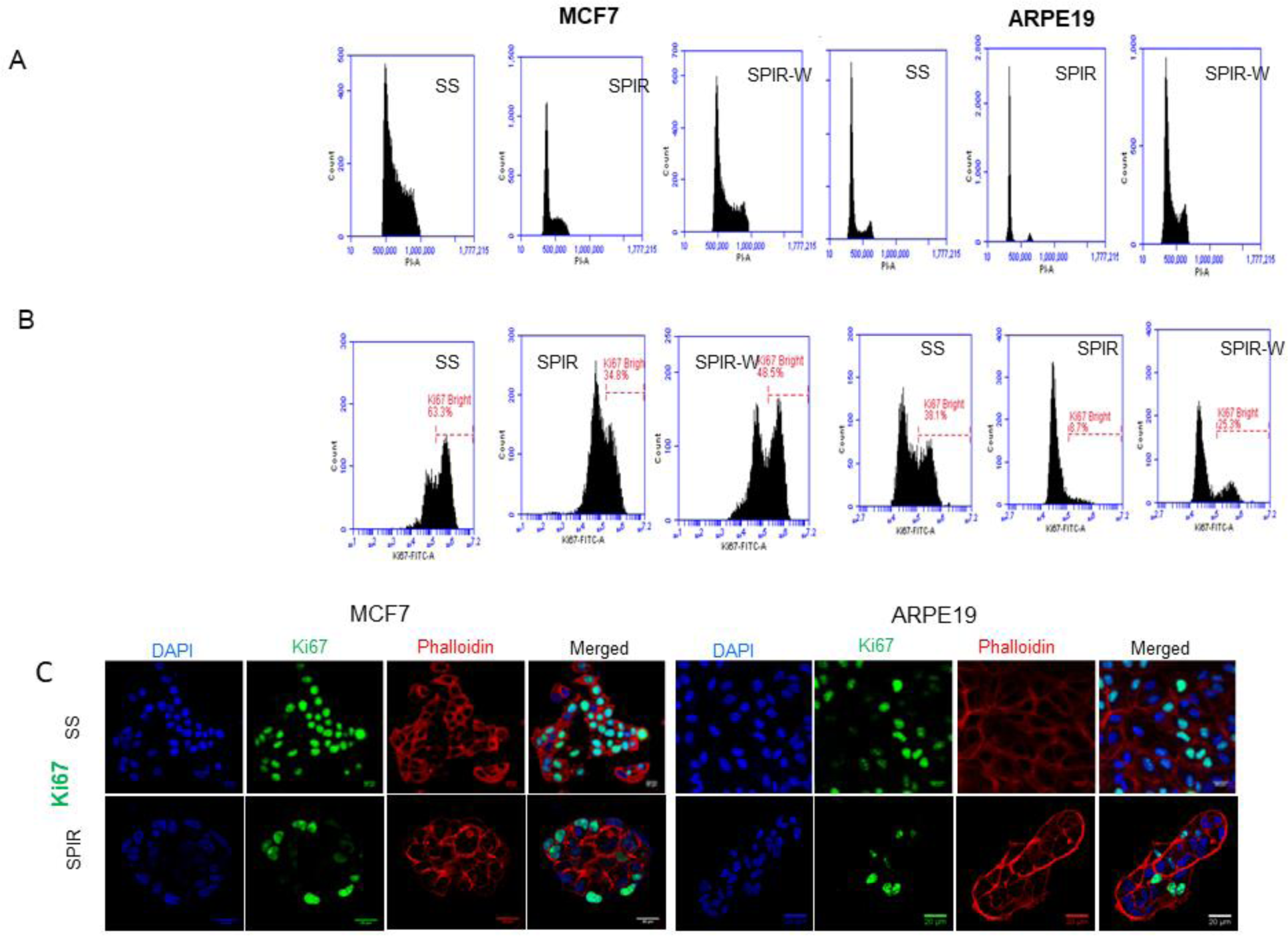
Panel A: Cell cycle analysis of MCF7 and ARPE-19 cells by flow cytometry. The initial and last peaks represent the G0-G1 and G2-M stage respectively; middle is S stage. Panels B and C show estimation of Ki 67 positivity by flow cytometry (B) and localization of Ki 67 by confocal microscopy (C) of MCF7 and ARPE-19 cells, under SS and SPIR conditions, as well as when trypsin has been removed from the culture following SPIR treatment (scale bar = 20 μm).

**Fig S9:**
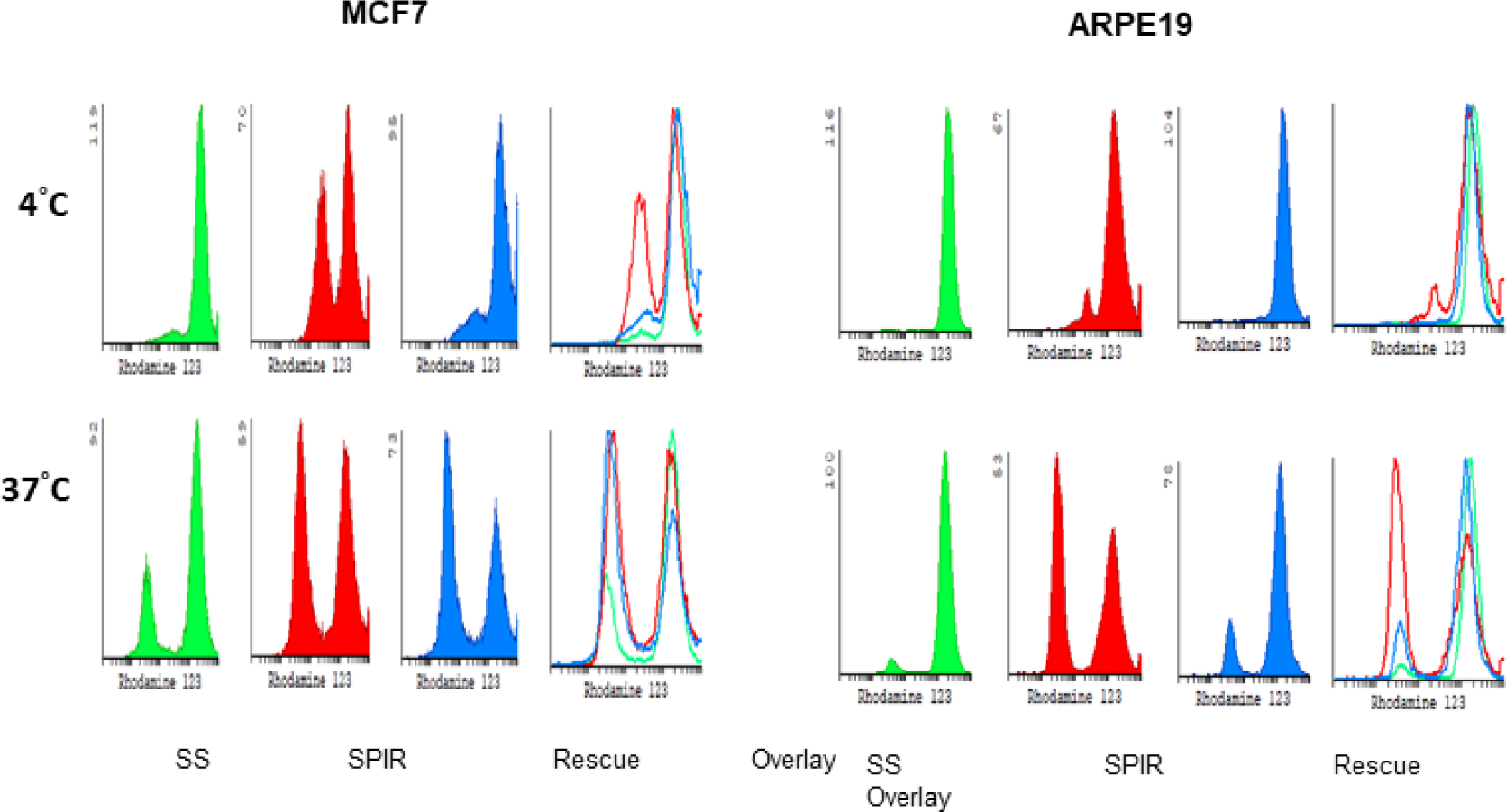
Flow cytometry histograms of rhodamine 123 uptake and efflux under SPIR conditions, SS conditions as well as when trypsin is removed from cell culture medium following SPIR treatment (SPIR-W).

**Fig S10:**
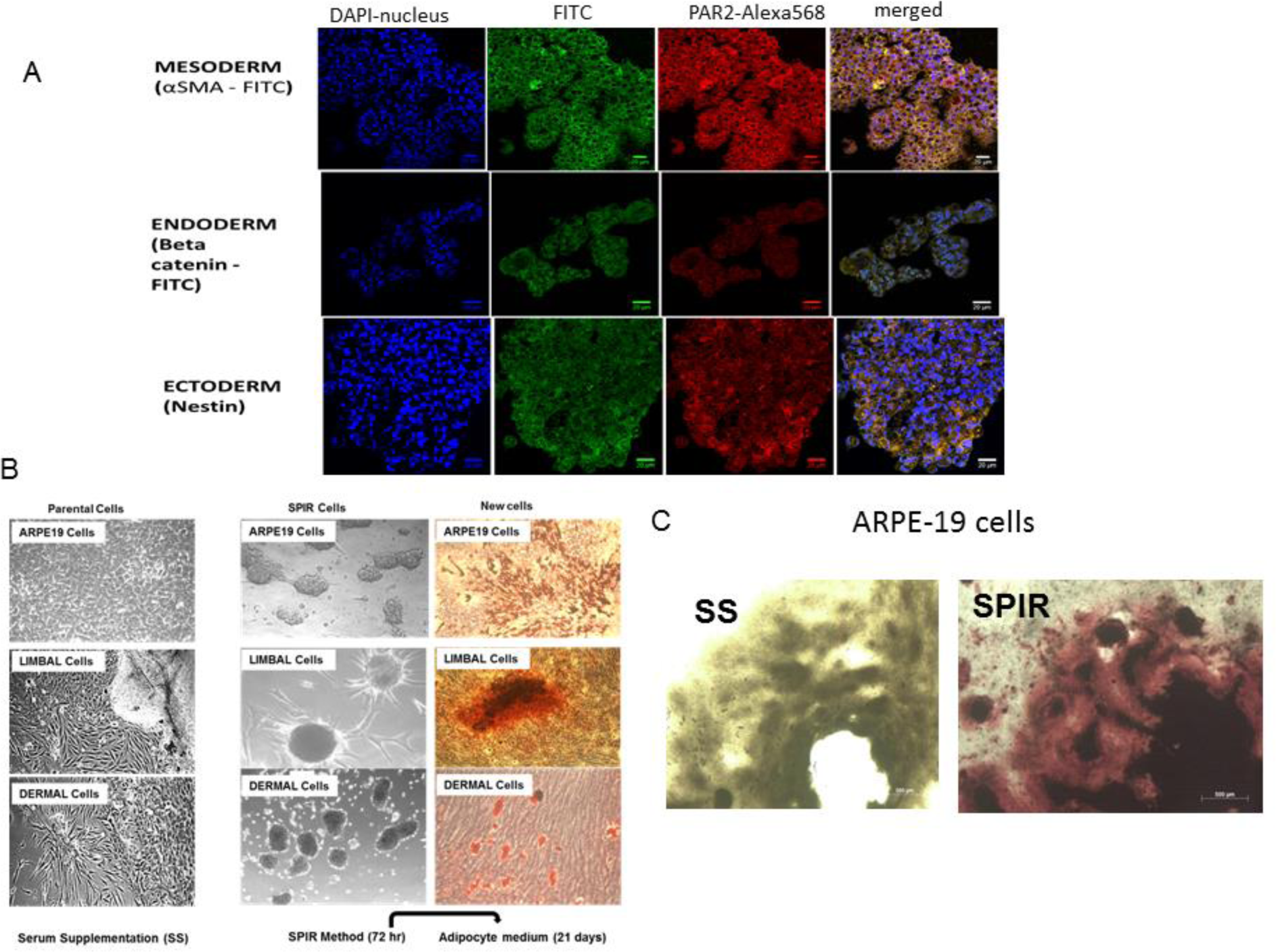
Panel A: Factor-dependent transdifferentiation of ARPE-19 cells into all three germ cell lineages enabled by SPIR treatment; Panel B: Adipogenic differentiation potential of ARPE19, limbal cells derived from corneal tissue (HCFs) and dermal fibroblasts, grown under SF conditions and differentiated using adipocyte culture medium; presence of oil droplets was determined by oil red-o staining; Panel C: Osteocytic differentiation of SPIR-treated ARPE-19 cells, determined by alizarin red staining.

**Fig S11:**
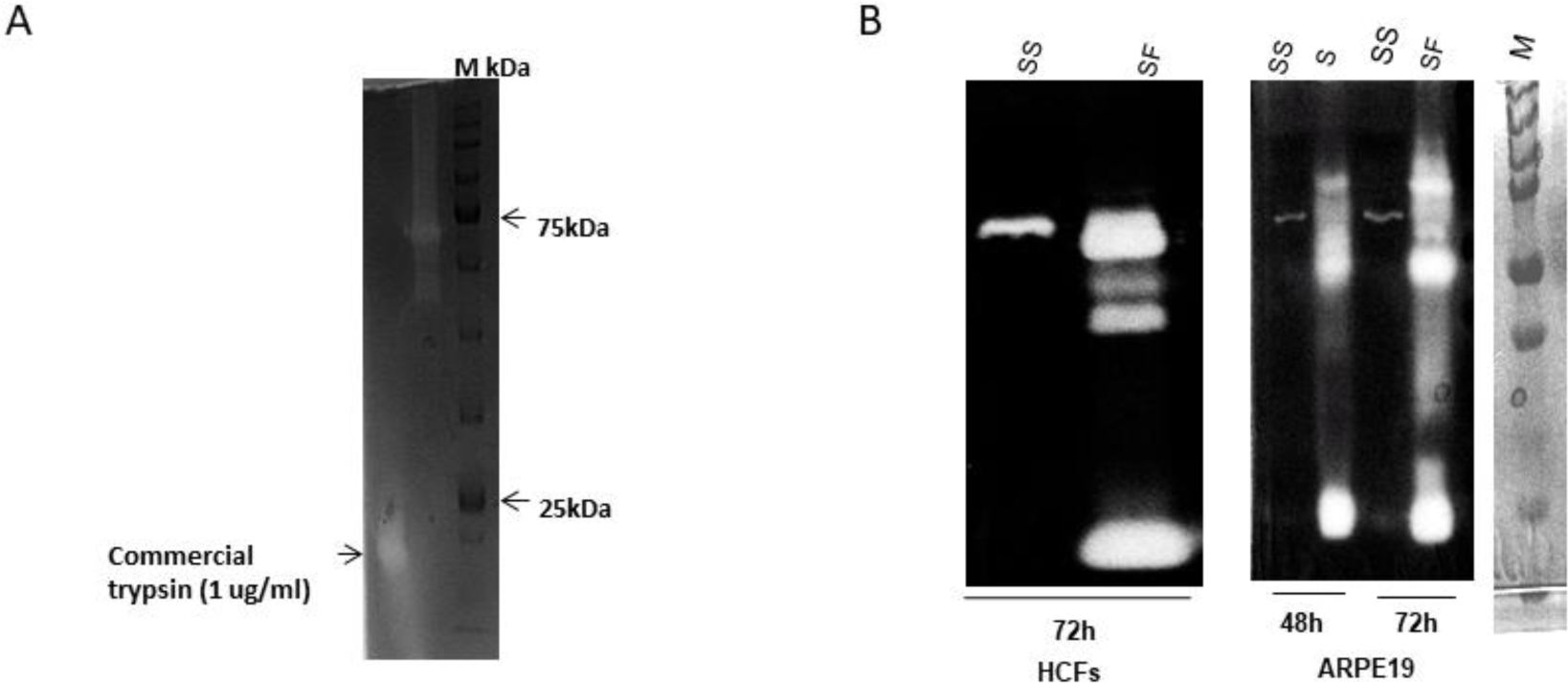
Panel A: Gelatin zymogram shows gelatinolytic activity of commercial trypsin (lane 1) migrating at 20 kDa; molecular weight marker (lane2); Panel B: Gelatin zymograms of Human corneal fibroblasts (HCFs) and retinal pigment epithelial cells (ARPE-19). CM of cells cultured under SF conditions displayed increased activation of pro-MMP −2, −9 to their active (MMP-2 and −9) forms and a unique/novel LMW band.

**Supplementary Table 2:** Patients’ characteristics (n=60)

|  |  |
| --- | --- |
| Mean Age (range) | 54 (19-81 years) |
| Mean tumour size (range) | 3.2 (1-6 cm) |
| Type of Surgery: |  |
| TMAC | 56 |
| Lumpectomy+ AC | 04 |
| Histology: |  |
| IDC, NST | 59 |
| Others | 01 |
| Histological Grade: |  |
| Grade 1 (%) | 01 (1.7%) |
| Grade2 (%) | 12 (20%) |
| Grade3 (%) | 47 (78.3%) |
| Ki67 index: |  |
| ≤14% (%) | 03 (5%) |
| >14% (%) | 57 (95%) |
| Axillary LN status: |  |
| Negative (%) | 30 (50%) |
| Positive (%) | 30 (50%) |
| PAR2 staining: |  |
| Negative (%) | 08 (13.3%) |
| 1+ (%) | 16 (26.7%) |
| 2+ (%) | 24 (40%) |
| 3+ (%) | 12 (20%) |

